# Designing Combinatorial Multiepitope Vaccine Candidates against Scrub Typus by a Proteome-wide Immunoinfomatics Approach

**DOI:** 10.1101/2024.03.05.583536

**Authors:** Swarna Shaw, Arka Bagchi, Debyani Ruj, Sudipta Paul Bhattacharya, Arunima Biswas, Arijit Bhattacharya

## Abstract

Scrub typhus (ST) is a mite-borne infection caused by the bacteria *Orientia tsutsugamushi* and a major public health concern in eastern and south eastern Asia. Early diagnosis and immediate intervention with antibiotics are required to limit the severity of the infection. Prophylaxis is compromised by the absence of an effective vaccine against *O. tsutsugamushi*. Development an effective vaccine is impeded due to extreme divergence of major surface antigens limiting cross-protectivity. Adopting a proteome-wide immunoinformatic approach, here, immunodominant outer membrane proteins are identified and conserved regions enriched in B-cell, Tc-cell, and Th-cell epitopes were mapped. Similarly, immunodominant stretches from salivary proteins of the vector were identified. Fifteen independent MEVs were designed by combining such segments and were analysed for physicochemical and immunological properties. Tertiary structure of the prioritized MEV was modelled, and interaction with TLR2, 4, and 5 was profiled through molecular docking and MD simulation. A glycosylation-deficient version of the MEV was also designed and analysed similarly. Both the MEVs were immune-simulated, which indicated elicitation of effective immune response. *In silico* cloning and expression were performed for bacterial and mammalian expression systems. Though the proteome-wide combinatorial approach with immunodominant segments demonstrated prospect, experimental validation for its efficacy against ST is warranted.

## Introduction

The mite-borne infection, Scrub typhus (ST) or tsutsugamushi disease, is caused by the Gram-negative α-proteobacterium *Orientia tsutsugamushi* (*Ot*) transmitted through the bite of an infected larval Trombiculidae mite (Chigger) ^1^. An estimated 1 billion people are at risk of the acute febrile ailment, and about 1 million new cases are reported each year in the Southeast and Asian-Pacific region^2^. The disease has been declared to be one of the most underdiagnosed/underreported infection, which often necessitates hospitalization, by the World Health Organization^3^.The disease is endemic and is emerging as a major public health concern in the region extending from Northern Japan and far Eastern Russia in the North, to Northern Australia in the South, and to discrete regions of India, Pakistan, and Afghanistan in the east, often mentioned as the ‘tsutsugamsuhi triangle’ area^4^.

The vectors and reservoirs of *Ot*, trombicuid mites *Leptotrombidium deliense* and *L. akamushi*, mediate transovarial transmission of the bacteria^5^. Infected chiggers, raised from infected eggs, attaches to the skin of rodents and human to collect tissue fluid meal and inoculates the bacteria in the course through salivary secretions. *Ot* infects peripheral dendritic cells in skin, and infection results dermal and epidermal necrosis leading to formation of ulcer, which is the preliminary sign of infection. Subsequently *Ot* spread through lymphatic system and infect endothelial cells systemically ^6^. Typical symptoms like headache, fever, myalgia, generalized lymphadenopathy, transient hearing loss, and rash onset following the systemic infection. If untreated the infection might progress to cause severe ailments including respiratory distress, gastrointestinal haemorrhage, and renal failure^6^. Variation in severity and fatality have been recorded for outbreaks ranging from self-limiting to a more severe condition with a case fatality rate of 6% in absence of chemotherapeutic intervention and 1.5% with treatment. The mortality rate rises up to 13% if standard treatment regime fails^7^. However such observations are yet to get correlated with genomic variations among strains. To prevent severe complications, early diagnosis and immediate chemotherapeutic intervention are exigent. The drug most commonly used are doxycycline, tetracycline, chloramphenicol, and azithromycin^8^. Strains that are resistant to doxycycline and chloramphenicol have been reported in northern Thailand and in China^9^. Despite consistent attemptsspanning several decades, an effective vaccine development is still awaited. Inactivated (formalin killed) *Ot* strains, formalin-fixed infected tissues, less-virulent strains, administration of virulent strain followed by antibiotic treatment, live irradiated *Ot*, subunit vaccines^10^, and DNA vaccine^11^have been attempted in this course for the purpose of developing an effective vaccine. The majority of the trials conferred protection for a period of <1yr. Moreover, the immune-memory developed by such vaccine trials were poorly cross-protective across diverse strains. More than 20 strains have been identified for *Ot* so far in addition to the type strains Gilliam, Karp, and Kato^4^. Though potentially immunodominant, tissue specific antigen (TSA56), a 56 kDa is an immunodominant surface antigen that demonstrates extreme heterogeneity which restricts cross-protectivity against divergent strains. Hence identification immunodominant as well as conserved *Ot*-specific vaccine candidates is exigent. In recent years immunization in mice with a combination ScaA, an autotransporter of *Ot*, and TSA56, a membrane protein demonstrated augmentation of protective immunity ^12^, suggesting identifying immunodominant regions from outermembrane proteins and configuring a multiepitope vaccine (MEV) might be a viable strategy in designing novel vaccines for ST. With the advances in immunoinformatics in developing a plethora of tools to predict epitopes, immunogenicity, allergenicity, and simulate possible immune responses against candidate vaccines, along with the generation of huge amount of genome-wide data, designing and configuring potential MEVs is gaining considerable attention as a reliable and rapid approach aiming identification of potential vaccine candidates^13^. Apart from aiming viral and extracellular bacterial pathogens, a number of potential candidates against intracellular bacterial pathogens like *Chlamydia*^14^ and *Brucella*^15^ have been identified.

One major impediment for vaccine development against *Ot* is the scarcity of comprehensive study to explore the immunocomponents for protection against *Ot* infection. Hauptmann et al., (2016) and Xu et al., (2017) demonstrated that adaptive transfer of CD8+ cells from *Ot*-recovered mice elicited robust protection ^16,17^. Kodama et al., (1988) reported protection or partial protection against various *Ot* strains by adaptive transfer of IFN-γ producing CD4^+^ cells^18^. Such results suggested that CD4^+^ protection is strain-dependent. Similarly, the passive transfer of primed serum against a specific strain was demonstrated to be ineffective towards infection protection against other strains ^19^. However, identifying conserved immunodominant antigens and combining multiple such epitopes to raise long-lived cross-reactive immunity might offer crucial clues for developing effective MEV. TSA56, reported earlier to potentiate protective immune response ^12^, has been targeted to develop a chimeric MEV candidate recently ^20^. Nano particle containing a chimeric vaccine candidate derived of the TSA56 and ScaA have been demonstrated to induce Ag-specific humoral immunity and T cell response ^21^. However other surface proteins of the pathogen remains unexplored in terms of immunogenicity or vaccine designing.

In this backdrop, a genome-wide approach was adopted to identify major outermembrane (OM) proteins of *Ot*. From the 2364 annotated proteins, 27 OM proteins were ranked for immunogenicity and scanned for immunodominant regions comprising multiple highly ranked B-cell, Th-cell, and Tc-cell epitopes. Two such segments were linked in combination to generate a set of immunodominant polypeptides comprising less than 300 amino acid residues. The saliva of haematophagous arthropods harbour a number of constituents including proteins with potential immunomodulatory action that are linked directly to infection establishment^22^. Saliva of *Ixodes scapularis*, the tick transmitting Lyme disease, triggers proinflammatory cytokine release by macrophages by impairing Toll-like (TLR) and Nod-like receptor (NLR) signalling pathways^23^. Humoral immunity directed against salivary proteins elicits acquired tick resistance and tick rejection^23^. Recently, a candidate vaccine comprising nucleoside-modified mRNAs encoding 19 *I. scapularis* salivary proteins (19ISP) has been demonstrated to confer resistance against the tick and restrained transmission of the pathogen in Guinea pigs^24^. Orthologues of *I. scapularis* salivary proteins in trombidid mite genomes have been annotated and enlisted ^25,26^. Considering combining eliciting memory against the pathogen and conferring tick resistance to mitigate transmission, here immunodominant epitopes from *L. deliense* salivary proteome are linked with the identified immunodominant stretches of OM protein of *Ot* to configure candidate MEV. Collectively 15 different MEVs were designed and each was evaluated for physicochemical properties, immunogenicity, allergenicity to prioritize a candidate MEV. The identified MEV (STMEV3) was characterized for structural and immunogenic properties which demonstrated it to be a stable vaccine candidate with the potential for profound immune-activation and memory development. A N-glycosylation deficient version of STMEV3 was generated by substituting specific Asn residues to Gln (STMEV3E). Three constructs were designed for expression of STMEV3 and STMEV3E. Molecular docking and simulation studies indicated stable interaction with TLR-4 and TLR-2. Taken together the data projects STMEV3 and STMEV3E as prospective vaccine candidates against ST.

## Methodology

### Sequence retrieval and prediction of subcellular localization

In the annotated genome *Orientia tsutsugamushi* str. Boryong (Genome ID: 357244.4) available at BVBRC, 2364 proteins are annotated. The sequences of the proteins were retrieved from Bacterial and Viral Bioinformatics Resource Center (BV-BRC) and the subcellular localization were analysed by PSORTb version 3.0.3 (RRID: SCR_007038) ^26^ for Gram negative bacteria. Product ID associated with the annotation is used as identifier of the proteins. Sequences *Leptotrombidium deliense* salivary proteins as enlisted by Dong et al. ^25^, were retrieved from Uniprot.

### Immunogenicuty and allergenicity prediction

Immunogenicity of proteins and configured MEVs were performed with VaxiJen 2.0 (RRID:SCR_018514) ^27^ the immunogenic proteins were raked based on the predicted *Protective Antigen Prediction* score. The MEVs were also profiled by ANTIGENpro (RRID:SCR_018779) ^28^ for predicting probability of antigenicity. Allergenicity of the antigens were predicted by AlgPred2.0 (RRID:SCR_018780)^29^ with a threshold of 0.3 for AAC based RF for allergenicity, and AllerCatPro2.0 ^30^. Toxicity of the MEVs were predicted through CSM-Toxin^31^ using default parameters.

### Prediction of sequential B-cell epitopes

Sequential B-cell epitopes on the OM-proteins were mapped using Bepipred Linear Epitope Prediction 2.0 and Bepipred Linear Epitope Prediction (RRID: SCR_018499). Epitopes with a length between 20-40 amino acids were prioritized and enlisted.

### Cytotoxic T-lymphocyte (Tc) and helper T-lymphocyte (Th) Epitopes

Using TepiTool hosted in IEDB server (RRID:SCR_006604) Tc-cell epitopes were predicted for 27 most frequent HLA-A and HLA-B alleles applying default settings for low number of peptides with IEDB recommended selection based on predicted consensus percentile rank. Th-cell epitopes were predicted were also predicted with TepiTool using the panel of 26 most frequent alleles for promiscuous binding applying default settings for moderate number of peptides with IEDB recommended selection based on predicted consensus percentile rank. For both the groups of epitopes lowest percentile rank represent a high affinity for MHC.

### Mapping immunodominant stretches and conservancy analysis

Nonoverlapping B-cell, Th-cell, and Tc-cell epitopes were mapped on each protein. 100-150 residue segments where at least two linear B-cell epitopes, three high raked (percentile rank ≤0.001) nonoverlapping MHC-II epitopes, and three high ranked (percentile rank ≤0.01) nonoverlapping MHC-I epitopes could be mapped were considered as immune dominant stretch on each protein. Six such stretches from the immunodominant OM proteins were enlisted and were analysed for conservancy across the ten *Ot* genomes based on bit-scores projected by BLASTP (RRID:SCR_001010). A hierarchical clustering were performed with heat map generation using Heatmapper (RRID: SCR_016974, http://heatmapper.ca/) generate a comprehensive view of the conservancy analysis. BlastP was also performed to compare the sequences of the segments with *Homo sapiens*.

### Analyzing physicochemical property

Physicochemical parameters like pI, instability index, aliphatic index, grand average of hydropathicity (GRAVY) etc. were estimated in ProtParam server (RRID: SCR_018087). Overall solubility of MEVs were estimated by Protein-Sol implementing default parameters. N-linked and O-linked glycosylation sites for bacterial proteins were identified by GLYCOPP (V 1.0) with a cut-off score of 0.5. After predicting the structures of MEVs, propensity of aggregate formation were evaluated by identifying possible aggregation inducing patches by AGGRESCAN (RRID: SCR_008403) using default parameters.

### Population coverage

Effectiveness across world population was predicted for the MEVs were performed by Population Coverage analysis hosted in IEDB for the predicted Th-cell and Tc-cell epitopes across most frequent MHC Class I and MHC Class II alleles as enlisted earlier^32,33^. Projected population coverage (PC), average number of epitope hits / HLA combinations recognized by the population (average hit), and minimum number of epitope hits / HLA combinations recognized by 90% of the population (PC90) were considered analysis.

### Structure prediction

Secondary structures of the MEVs were predicted using PSIPRED 4.0. Tertiary structures of MEVs were predicted by RoseTTAFold (RRID: SCR_018805) hosted in Robetta^34^. The structures of the MEVs were also predicted by implementing ColabFold v1.5.3: AlphaFold2 (RRID: SCR_023662)^35^ using MMseqs2 with 8 seeds and 3 recycles. The top ranked models were refined using the Galaxy Refine (RRID: SCR_006281)^36^ server. The model quality for MEVs has been assessed by the ProSA-web server (RRID: SCR_018540, https://prosa.services.came.sbg.ac.at/prosa.php) for Z-score prediction ^37^. The Ramachandran plot was generated using the PROCHECK (RRID: SCR_019043, https://saves.mbi.ucla.edu/) server to generate the Ramachandran plot and enumerate residues in the disallowed region. Models with the least number of residues in the disallowed region were selected for following studies. The structures were visualized with PyMOL (RRID: SCR_000305).

### Conformational epitopes, IFN epitoes, Aggregation, glycosylation

To map conformational B-cell epitopes, Ellipro server^38^ was implemented to identify discontinuous B-cell epitopes on the predicted models for MEVs with a cut-off score of 0.5 and minimum distance of 6Å. The tool implements a combination of approaches to predict antigenicity of surface protrusion and predicts discontinuous epitopes based on protrusion index (PI) values. The epitopes were visualized with PyMOL. IFN-γ epitopes were predicted from the enlisted Th cell epitopes by IFNepitope (RRID: SCR_024745)^39^ in hybrid approach (machine learning technique and motifs-based search).

### Codon optimization and in silico cloning

Codon optimizations for *E. coli* K12 and Homo sapiens were performed while reverse translating the MEV amino acid sequences using EMBOSS Transeq (RRID: SCR_015647). SnapGene 5.1.5 (RRID: SCR_015052) software was implemented to clone the codon optimized sequence into *E. coli* pET-16b (+) vector. After the restriction sites of *Nde*I and *Bam*H1 were incorporated in the 5’ and 3’ -terminal sequences, respectively. Aiming expression in human, restriction sites for *Hin*dIII and *Xba*I sites were incorporated in the 5’ and 3’ respectively for the human codon optimized back translated DNA with restriction sites for *Hin*dIII and *Xba*I in the 5’ and 3’ end respectively. For optimal expression in mammalian expression system, CACC sequence was introduced as a part Kozak signature was at the upstream of the start codon. For generating a secretory protein expressing construct human somatotropin signal sequence ‘MATGSRTSLLLAFGLLCLPWLQEGSAFPTIPLS’ was fused with the MEV.

### Immune Response Simulation

Possible immunological responses of the MEVs were projected using the dynamic immune simulation server C-ImmSim online (RRID: SCR_018775)^40^ for three injections at 30 days interval. The simulation was performed with a simulation volume of 10 and 1000 steps for default MHC allele set (A0101, A0101, B0702, B0702, DRB1_0101, and DRB1_0101).

### Protein-protein docking

Protein-protein docking was performed independently with the predicted structures of MEVs and the available structures for TLR2 (3A79, chain A), TLR4 (3VQ2, Chain A, B), and TLR5 (3J0A, Chain A) in PDB by Hdock (RRID: SCR_024799)^41^ implementing default parameters. The complexes were visualized with PyMOL. The highest ranked docked models were analysed for predicting binding energy and K_d_ by PRODIGY ^42^. The interactive residues contact maps were generated using COCOMAPS ^43^.

### MD-simulation

The Molecular dynamics (MD) simulations were performed using GROMACS (RRID:SCR_014565)^44^ software packages (version 2022.4 and 2022.3) in a terascale high performance Linux (CentOS 7.9) based computing cluster at the S. N. Bose Innovation Centre of University of Kalyani, Kalyani, Nadia, West Bengal, India. The coordinates (in .gro file format), topology (in .top file format) and the position restraints (in .itp file format) of the proteins were created from the Protein Data Bank (PDB) format using ‘*pdb2gmx’* function of GROMACS after applying Charmm27 force field. TIP3P water model was used to solvate the proteins and the total charge of the solvated systems were neutralized with Na^+^ and Cl^-^ ions (at a concentration of 0.1M). The energy minimization of the system was carried out using steepest descent algorithm in GROMACS with maximum number of steps set to 50000 and until a maximum force of <10kJ/mol was obtained. The thermodynamic equilibration of the system was carried out using NVT and NPT ensembles available in GROMACS. During NVT equilibration, V-rescale thermostat, a modified Berendsen thermostat, was used for temperature coupling. On the other hand, C-rescale barostat was utilized for pressure coupling during NPT equilibration. Once the configuration files were ready, trajectory files have been generated by GROMACS, which described the phase space dynamics of the atoms of the complexes oer a period of 100 nanoseconds (ns). The root-mean-square deviation (RMSD), root-mean-square fluctuation (RMSF), radius of gyration (Rg)of the proteins and the number of stable hydrogen bonds between the proteins of the docked complexes were analysed with the help of tools internal to GROMACS and the output files were plotted using Matplotlib (RRID: SCR_008624) and NumPy (RRID: SCR_008633) to obtain line graphs.

## Results

### Mapping immunodominant regions in the outer membrane proteins of Orientia tsutsugamushi

A total of 2364 protein sequences from *Orientia tsutsugamushi* (str. Boryong) are available in the Bacterial and Viral Bioinformatics Resource Center (BV-BRC). Each of the protein sequences was analysed for subcellular localization in Gram-negative bacteria by PSRTb. The analysis predicted localization of 826, 256, 7, 1190, and 27 proteins in the cytoplasm, cytoplasmic membrane, extracellular, periplasm, and outer membrane respectively while for 58 proteins localization could not be predicted (Fig. 1A). Considering surface exposure and accessibility, 27 outer membrane proteins were further explored for identifying immunodominant regions. A primary prediction for immunogenicity was performed for the proteins with VaxiJen2.0. As depicted in Fig. 1B, the proteins demonstrated an *Overall Protective Antigen Prediction* score between 0.2926 (for 357244.4.peg.1590) to 0.7326 (for 357244.4.peg.423). Considering a cut-off of 0.5 for the *Overall Protective Antigen Prediction* score as an indicator for potential immunogenicity, a total of 7 proteins comprising 357244.4.peg.423 (score: 0.7326), 357244.4.peg.643 (score: 0.6835), 357244.4.peg.1197 (score: 0.6168), 357244.4.peg.678 (score: 0.6168), 357244.4.peg.2076 (score: 0.5668), 357244.4.peg.625 (score: 0.5665), and 357244.4.peg.15 (score: 0.5463) were identified as potentially immunogenic outer membrane proteins. Since the sequences of 357244.4.peg.1197 and 357244.4.peg.678 are identical, effectively 6 proteins were prioritized for subsequent analysis.

**Fig. 1.**
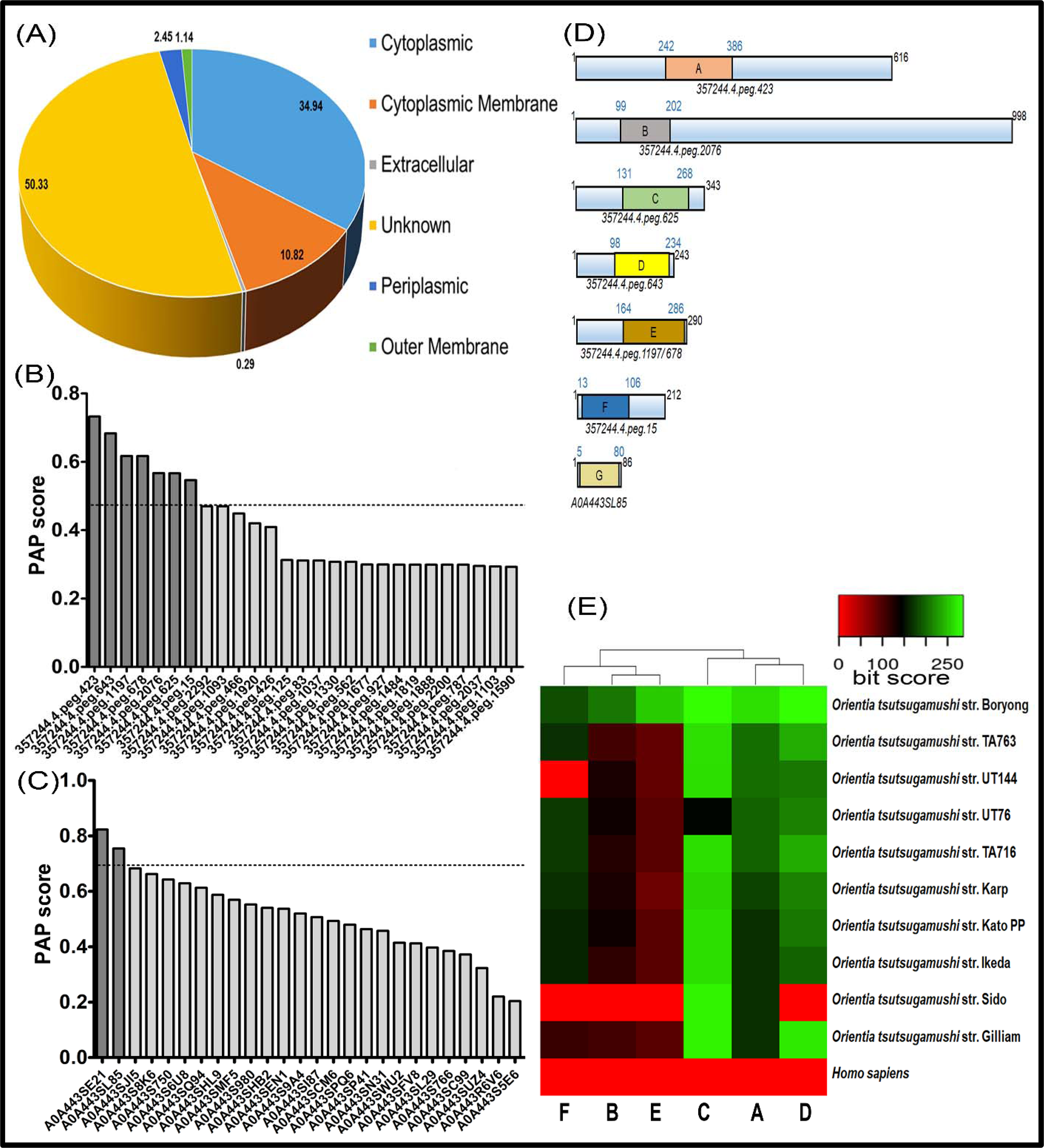
Proteome-wide identification of immunodominant proteins and immunogenic stretches. Amino acid sequences of 2364 proteins from *O. tsutsugamushi* were analysed by *Psortb* for possible subcellular localization. The % of proteons in various subcellular localization are projected in the pie chart (A). 27 OM-proteins from *O. tsutsugamushi* (B) and 26 salivary proteins from *L. deliense* (C) were analysed for antigenicity by Vaxijen 2.0. (D) 75-150 amino acid long stretches enriched in B-cell, Th-cell, and Tc-cell epitopes were mapped on the OM proteins outer membrane proteins of *O. tsutsugamushi* and salivary protein of *L. deliense*. Heat map presentation and hierarchical clustering of immunodominant stretches based on Bitscore obtained for the hits from the strains while performing BALSTP (E).

Sequential B-cell epitopes were predicted from each of the six prioritized proteins implementing BepiPred and BepiPred-2.0 (Table-S1). T-cell epitopes for class I and class II MHCs were identified by TepiTools for 27 most frequent HLA-A and HLA-B alleles, and 26 most frequent HLA-DP, HLA-DQ, and HLA-DR alleles respectively. Less than 0.2-percentile ranked 9-mer high-binding MHC-I epitopes (Table-S2) and 15-mer high-binding MHC-II epitopes (Table-S3) were enlisted from the prioritized 6 immunodominant outer membrane proteins. Potentially immunogenic regions from each of the proteins were identified by mapping 100-150 residue stretch featuring at least two linear B-cell epitopes, three high-raked (percentile rank ≤0.001) nonoverlapping MHC-II epitopes, and three high-ranked (percentile rank ≤0.01) nonoverlapping MHC-I epitopes (segment A: 357244.4.peg.423, L242 to S386; segment B: 357244.4.peg.2076, P99 to S202; segment C: 357244.4.peg.625, V131 to A268; segment D: 357244.4.peg.643, T98 to T234; segment E: 357244.4.peg.1197, A164 to V286; and segment F: 357244.4.peg.15, Y13 to S106) (Fig. 1D). The identified regions from each of the proteins along with the epitopes identified are elaborated in Table-1 and the work-flow for identifying such segments is presented in Fig. S1. Conservancy analysis of each of the selected segments was performed by BLASTP against *Ot* (taxid: 784). As presented in Table-S4, two of the 6 selected segments resulted in at least one hit against the data set available for 10 major strains of *Ot* and the rest demonstrated at least one hit against nine strains. When investigated for possible similarity with any human proteins by performing BLASTP against *Homo sapiens* (taxid 9606), none of the selected segments offered a hit indicating the absence of similar sequences in human proteome (Table-S4). A hierarchical clustering was performed based on bit scores obtained from the BASTP analysis for each segment against ten *Ot* strains. As represented through the heatmap in Fig. 1E, segments C, A, and D clustered with higher bit-scores for the majority of the strains, suggesting greater conservancy for the segments.

**Table1:**
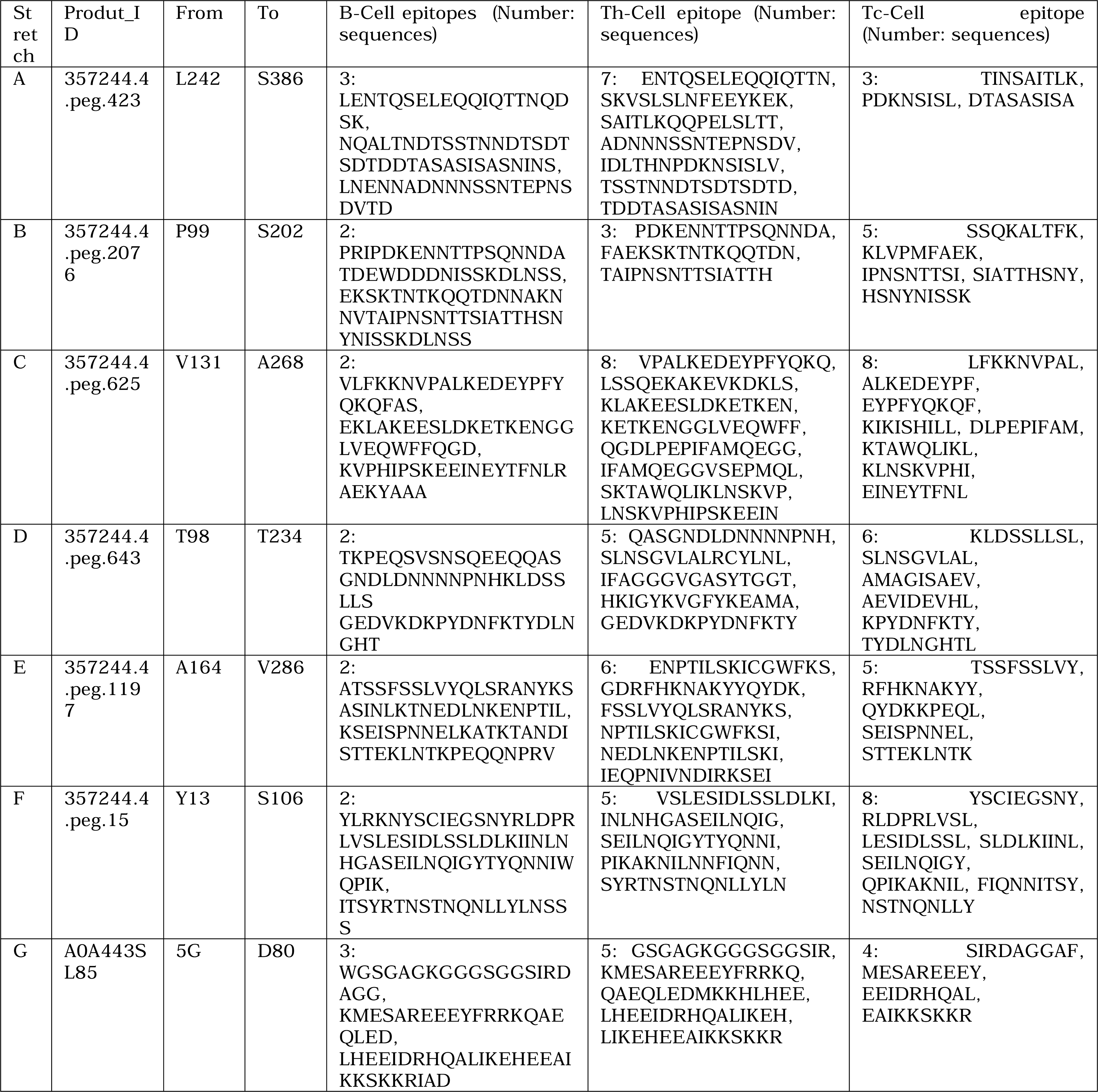
Detail of the immunodominant stretches and list of identified epitopes. Selected stretches enriched in B-cell, Tc-cell, and Th-cell epitopes from immunodominantoutermembrane proteins from *Ot* and salivary protein of *L. deliense* are enlisted with details of number of B-cell, Tc-cell, and Th-cell epitopes and the sequences.

### Mapping immunodominant regions in the salivary proteins of Leptotrombidiumdeliense

Tick salivary proteins play a crucial role in vector borne infections. Such proteins have also been exploited in vaccine designing and raising protective immune-memory ^24^). From a list provided by Dong et al., ^25^ for salivary proteins in trombidid mite genomes, 26 protein sequences of salivary proteins of *Ixodes scapularis* were retrieved from the Uniprot database. Implementing BLASTP, orthologues for each of the salivary proteins were identified from the available protein sequences from *L. deliense* (299467). The proteins were analysed for immunogenicity by VaxiJen2.0 which identified large ribosomal subunit protein P2 (A0A443SE21, *Overall Protective Antigen Prediction* score: 0.8231) and ATPase inhibitor-like protein (A0A443SL85, *Overall Protective Antigen Prediction* score: 0.7546) as potentially immunogenic (Fig.1C). Since A0A443SE21 demonstrated 64% identity and 81% similarity with the human orthologue (2LBF_B, Fig. S2A), A0A443SL85 was selected for identification of the immunodominant segment. Sequential B-cell epitopes were predicted by BepiPred and BepiPred-2.0 (Table-S1). Four high-ranked T-cell epitopes for class I (percentile rank ≤0.01) and five class II (percentile rank ≤0.001) MHCs were identified by TepiTools for the 27 most frequent HLA-A and HLA-B alleles, and the 26 most frequent HLA-DP, HLA-DQ, and HLA-DR alleles respectively. Less than 0.2-percentile ranked 9-mer high-binding MHC-I epitopes (Table-S2) and 15-mer high-binding MHC-II epitopes were enlisted from the protein (Table-S3). Potentially immunogenic regions from each of the proteins were identified by mapping a 75 residue stretch (between G5 to D80, mentioned as segment G from here) featuring several linear B-cell epitopes, MHC-II epitopes, and MHC-I epitopes (Table-1, Fig. 1D, and Fig. S1). The region demonstrated mere 50.79% identity with the human orthologue (Fig. S2B).

### Configuring multiepitope vaccine candidates and mapping conformational B-cell epitopes

Following the identification of potentially immunogenic regions if the OM proteins (segment A to F) of *Ot* and salivary proteins of *L. deliense* (segment G), prospective multiepitope vaccines (MEV) was configured by combining two such segments from OM proteins with segment G. Collectively, 15 such combinations could be generated (Fig. 2A). Since the selected regions were flanked by sequential B-cell epitopes, the segments were linked by a KK linker as described earlier by Kumar et al. ^45^. Each of the combinations were considered as candidate multiepitope vaccine (STMEV1 to 15) and were analysed for immunogenicity. As presented in Fig. 2B, STMEV3 demonstrated the highest immunogenicity score (0.9156) followed by STMEV1 (0.8911), STMEV5 (0.8764), STMEV7 (0.8678), and STMEV4 (0.8416). Similar analysis with ANTIGENpro predicted highest antigenicity of 0.952 for STMEV3, followed by STMEV7 (0.946) (Fig. 2B). Predicted allergenicity of the MEVs were estimated with AlgPred2.0 with RF model and with AllerCatPro2.0. While AlgPred2.0 assigned STMEV10, STMEV13, and STMEV7 as allergens with ML score of 0.427, 0.328, and 0.302 respectively (Fig. 2B), none of the MEVs were assigned as allergen by AllerCatPro2.0 (Table-S5). Similarly, toxicity prediction with CSM-Toxin projected none of the MEVs to be toxic. Physicochemical properties of the MEVs were analysed with ProtParam. Theoretical pI was determined between 4.84 to 9.36 for the 15 MEVs, while the aliphatic index ranged between 60.27 to 80. The Grand average of hydropathicity (GRAVY) for the MEVs ranged between -1.137 to 0.708 (Table-2). Though the majority of the MEVs were analysed to be stable, STMEV12, STMEV6, STMEV11, STMEV10, STMEV15, and STMEV9 were predicted to be unstable with the instability index above 40. When solubility of the MEVs were predicted by Protein-Sol, STMEV1, STMEV5, STMEV3, STMEV2, and STMEV8 demonstrated maximum solubility with a predicted scaled solubility above 0.8 (Table-2). Toxicity prediction with CSM-Toxin projected none of the MEVs to be toxic. Collectively, considering immunogenicity, allergenicity, toxicity, and predicted physicochemical properties including stability, STMEV3 was prioritized for further exploration as candidate MEV. Population coverage of the STMEV3 was calculated with the *Population Coverage* tool hosted in IEDB server. Analysis for both class-I and class-II MHC alleles were performed with 22 and 27 most frequent HLA alleles respectively for the world population. As represented in Figs. 2C and 2D, for class-I MHC, STMEV3 demonstrated coverage, average hit, and pc90 of respectively. Similar analysis for the MEVs for class-II MHC indicated coverage, average hit, and pc90 of 98.55%, 29.88, and 19.69 respectively (Figs. 2C and 2E). The data suggested satisfactory population coverage for the MEV.

**Fig. 2.**
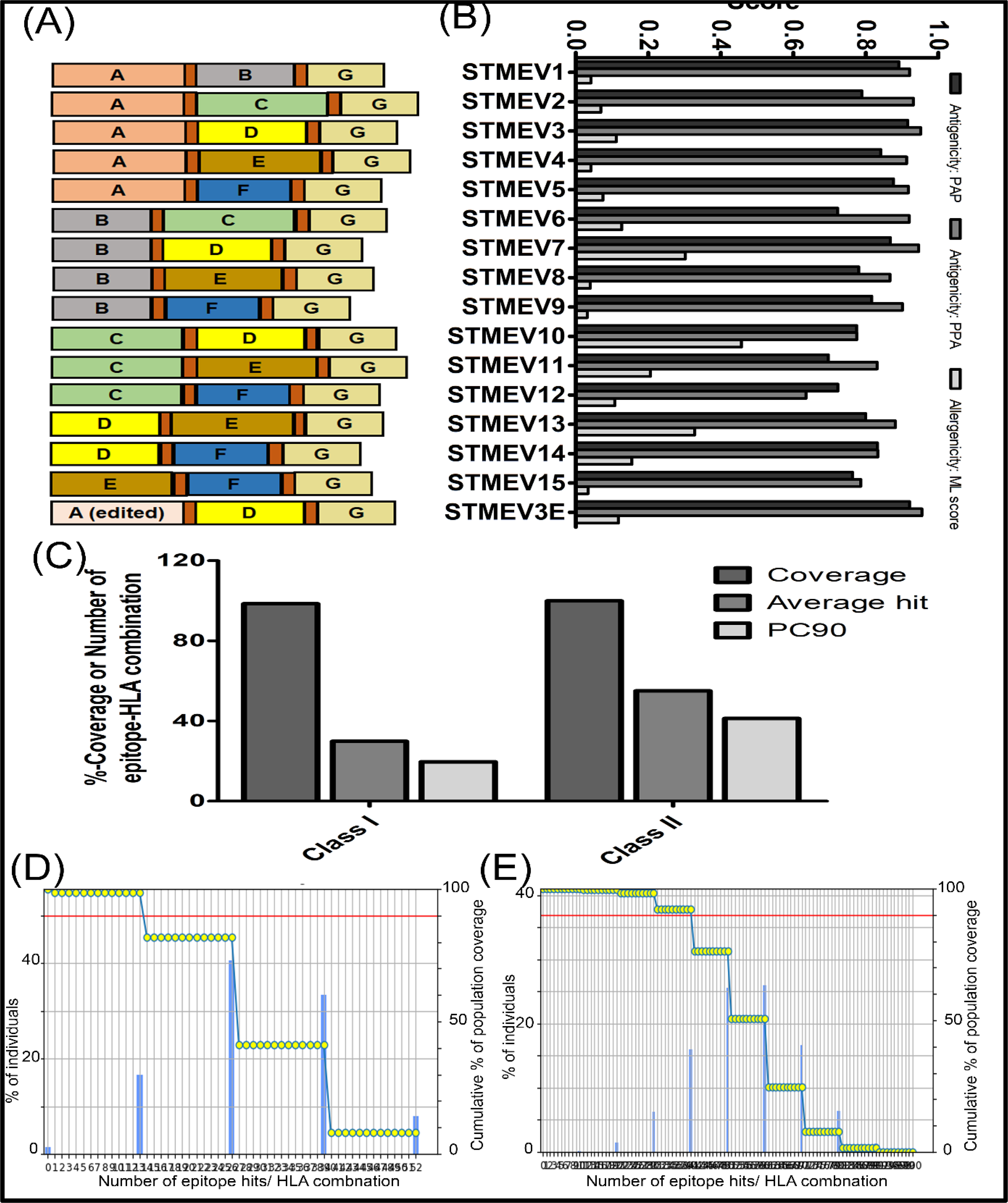
Multiepitope vaccine construction and population coverage analysis. Fifteen individual MEVs (STMEV 1 -15) were designed by combining immuodominant stretches [A-F regions from Fig. 1D] of outer membrane proteins of *O. tsutsugamushi* and immunodominant stretch [G region from Fig. 1D] from salivary protein of *L. deliense*. The configuration of STMEV3E is also included. (F) Antigenicity was predicted based on the PAP score as determined by Vaxijen 2.0 and PPA score calculated by AntigenPro. Allergenicity was predicted based on the ML-score enumerated by AlgPred2.0 (G). (H) Coverage (%), average hit, and PC90 for class I and class II MHC epitopes of STMEV3. Number of epitopes vs. % of individuals or cumulative % of population coverage plot for class I (I) and class II (J) MHC epitopes from STMEV3.

**Table 2.**
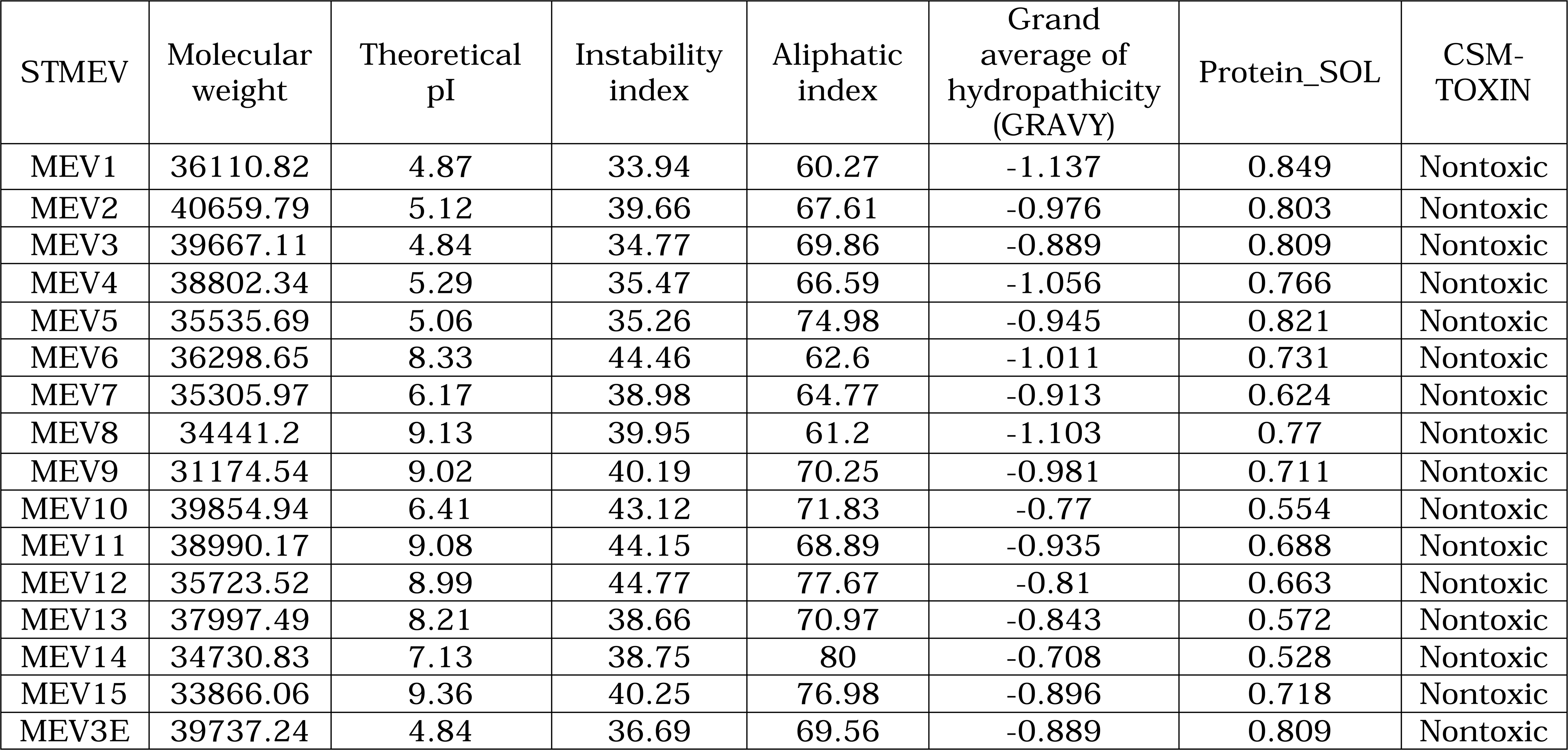
Physicochemical properties of the configured MEVs. Molecular weight, theoretical pI, instability index, aliphatic index, and grand average of hydropathicity (GRAVY) were determined by PROTPARAM. Protein solubility and toxicity were predicted by Protein-Sol and CSM-Toxin server respectively.

Glycosylation of proteins often eventuates glycan masking of epitopes on immunogens, which sequester the immunogenic surface from humoral immune surveillance strategies including recognition by B-cell receptors ^46^. Since the designed MEVs were constituted from prospective immunodominant regions, the absence of glycosylation sites would enhance their efficacy as vaccine candidates. STMEV3 was analysed for bacterial N-linked and O-linked glycosylation sites by GLYCOPP (V 1.0). As presented in Fig. 3A, though no O-lined glycosylation sites were predicted, five potential N-linked glycosylation sites were predicted at N78, N79, N116, N123, and N143. In order to mitigate the possibility of N-linked glycosylation without perturbing the physicochemical property, each of the predicted N (Asn) residues was substituted with Q (Gln) to generate STMEV3^N78Q, N79Q, N116Q, N123Q, N143Q^ (edited STMEV3, referred to as SETMEV3E here on). Analysing physicochemical properties indicated a theoretical pI of 4.84, aliphatic index of 69.56, and GRAVY of -0.889 (Table-2). With an instability index of 36.69 and scaled solubility of 0.809 the STMEV3E was predicted to be stable and soluble (Table-2). The MEV demonstrated immunogenicity score of 0.9206 (Vaxijen 2.0) and 0.955 (AntigenPro), suggesting it as a potential immunogen (Fig. 2B). Allergenicity prediction by AlgPred2.0 indicated it as nonallergen with a ML score of 0.118 (Fig. 2B) and identical prediction was made by AllerCatPro2.0 (Table-S5). Alongside STMEV3, STMEV3E was also explored further as candidate MEV.

**Fig. 3.**
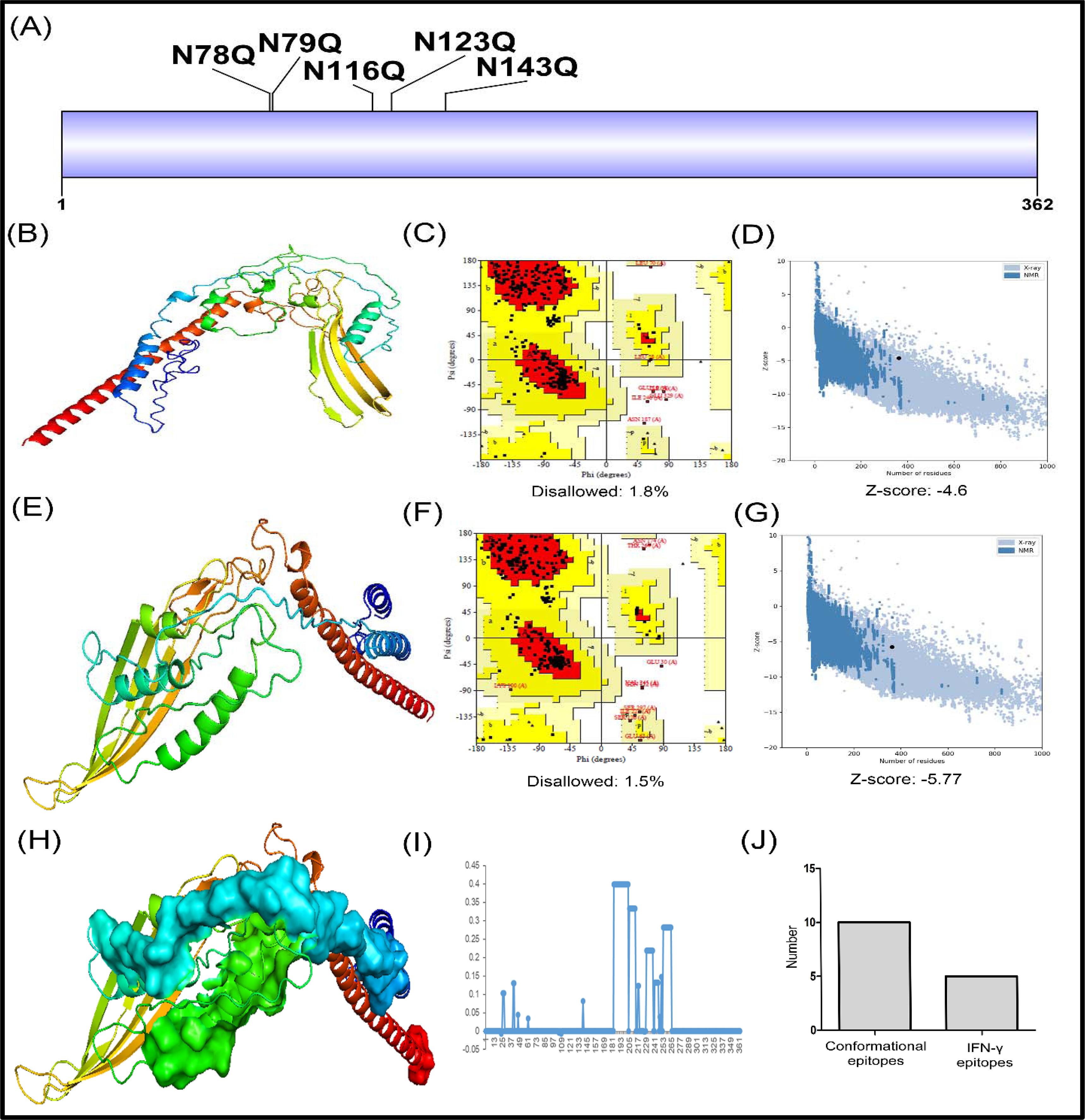
Glycosylation sites on STMEV3 and glycosylation deficient version of it (STMEVE), and structural characterization of STMEV3. (A) Probable N-glycosylation sites on STMEV3 were edited by Asn to Gln substitution to generate STMEV3E. (B) *Ribbon* diagram of the structure predicted for STMEV3E coloured in spectrum as predicted by RoseTTAFold and visualized by PyMol. Validation of the predicted structure by Ramachandran plot (C), percentage of residues in the disallowed region is mentioned separately; and by ProSA (D). Z-score is mentioned separately. (E) *Ribbon* diagram of the structure predicted for STMEV3 coloured in spectrum as predicted by RoseTTAFold and visualized by PyMol. Validation of the predicted structure by Ramachandran plot (F), percentage of residues in the disallowed region is mentioned separately; and by ProSA (G). Z-score is mentioned separately. (H) Major conformational B-cell epitopes predicted by Ellipro represented in *surface* view on the *ribbon* diagram of STMEV3. (I) Aggregation prone stretches as mapped by AGGRESCAN based on a4vAHS score of each residues. (J) Number of conformational B-cell epitopes and number of IFN-γ inducing Th-cell epitopes as predicted by IFN epitope.

### Structural analysis of STMEV3 and STMEV3E

The amino acid sequences of STMEV3 and STMEV3E were analysed for secondary structure with PsiPred. A number of internal alpha helices and beta strands were predicted along with two long terminal alpha helices in the MEVs (Figs. S3A and S3B). The tertiary structures of the MEVs were predicted with ColabFold (AlphaFold in Googlecolab; greedy pairing strategy, recycle 3 and seeds 8 and RoseTTAFold (Fig. 3B and 3E) in Robetta. The top-ranked model generated by ColabFold was refined by GalaxyRefine. The models were evaluated by assessing % of residues in the *disallowed* region in the Ramachandran Plot. While structure predictions for STMEV3 and STMEV3E generated by ColabFold demonstrated 1.5% and 1.2% of residues in the *disallowed* region, models generated by RoseTTAFold for the two MEVs demonstrated 1.5% and 1.6% residues in the *disallowed* region (Figs. 3F and 3C). The models were further verified by *ProSA* for predicting Z-score. While models for STMEV3 and STMEV3E generated by ColabFold demonstrated Z-score of -2.66 and -2.55, for models generated by RoseTTAFold Z-scores of -5.77 and -4.6 were determined respectively (Figs. 3G and 3D), indicating better over-all model quality for the RoseTTAFold predicted structures. Hence the RoseTTAFold-predicted structures were implemented in subsequent studies.

Conformational epitopes were mapped on the predicted structures with Ellipro. At least ten potential discontinuous epitopes could be mapped with four epitopes dispalying prediction score higher than 0.75 in STMEV3 involving 3 to 44 amino acids (Figs. 3H and 3J). For STMEV3E, seven discontinuous epitopes could be mapped with the prediction score higher than 0.75 in STMEV3E involving 4 to 30 residues (Table-S6). From the predicted structure possibility of aggregation for STMEV3 was evaluated by AGGRESCAN analysis. As shown in Fig. 3I, five potential aggregation prone regions could be predicted with modest AGGRESCAN score with in the range of 0.131 to 0.399.

### Codon optimization and in silico cloning of STMEV3 and STMEV3E

For *in silico* cloning and expression of *stmev3* and *stmev3e*, the amino acid sequences were back-translated using *Backtranslate*. *Backtranslate* was conducted with codon optimization setting for *E. coli* K12 and *Homo sapiens*. For bacterial expression, the *E. coli* K12 codon optimized fragment was cloned within *Nde*1 and *Bam*H1 sites of pET16b using SnapGene to generate the construct pET16bSTMEV3 and pET16bSTMEV3E. *In silico* translation of the construct resulted in to synthesis of STMEV3 and STMEV3E with N-terminal 6X-tag (Figs. S4A and 4A). For expression in mammalian cell two types of constructs were generated. The first construct was aimed at the production of a nonsecreted version of the MEVs. *Kozak* sequence (CACC) was added at upstream of the ORF for *stmev3* and *stmev3e*, codon optimized for *H. sapiens* (Figs. S4B and 4B). For the secretory version, the secretory signal ‘MATGSRTSLLLAFGLLCLPWLQEGSAFPTIPLS’ was added to the N-terminus of STMEV3 and STMEV3E. The resulting sequences were back-translated with codon optimized for *H. sapiens* and the *Kozak* sequence (CACC) was added at the upstream (Figs. S4C and 4C). The back-translated sequences were cloned in pcDNA3.1 within restriction sites for *Hind*III and *Xba*I. *In silico* translation of the constructs resulted in the synthesis of the target protein.

### STMEV3 and STMEV3E interaction with Toll-like receptors

Cell surface TLRs, particularly TLR2, TLR4, and TLR5, are involved in activating the antigen-presenting cells, proliferation of Th-cells, and augmenting IFN-γ and IL-2 production ^47,48^. To envision possible interaction of the TLRs with STMEV3 and STMEV4, structures of the TLR2 (3A79, chain A), TLR4 (3VQ2, Chain A, B), and TLR5 (3J0A, Chain A) were retrieved from PDB. The FFT-based global docking program integrated with available binding interface information, Hdock, was implemented to dock the TLRs with the predicted models of STMEV3 and STMEV3E independently. The contact point residue pairs were predicted by also predicted by Hdock (Table-S7).

The interactions were further explored by PRODIGY to estimate binding energy (ΔG) and dissociation constant (K_d_). STMEV3 was observed to interact with each of the TLRs with predicted binding energy of -11.8 kcal/ mol, -15 kcal/ mol, and -16 kcal/ mol and K_d_ of 2.3X10^-9^ M, 1.1X10^-11^ M, and 1.7X10^-12^ M for TLR2, TLR4, and TLR5 respectively (Figs. 5A, 5B, and 5C). For STMEV3E, the binding energieswere estimated to be -12.3 kcal/ mol, -11.3 kcal/ mol, and -18 kcal/ molwhile K_d_s were projected to be 9.8X10^-10^ M, 5.1X10^-9^ M, and 6.2X10^-14^ M for TLR2, TLR4, and TLR5 respectively (Figs. 5G, 5H, and 5I). 100, 125, and 74 residue pairs were predicted for STMEV3-TLR2, STMEV3-TLR4, and STMEV3-TLR5 interactions (Figs. 5D, 5E, and 5F, Table-S7). For STMEV3E, 79, 150, and 137residue pairs could be predicted while forming complexes with TLR2, TLR4, and TLR5 respectively (Figs. 5J, 5K, and 5L, Table-S7). Thus the molecular docking study indicated potential interaction between the TLRs and the MEVs.

**Fig. 4.**
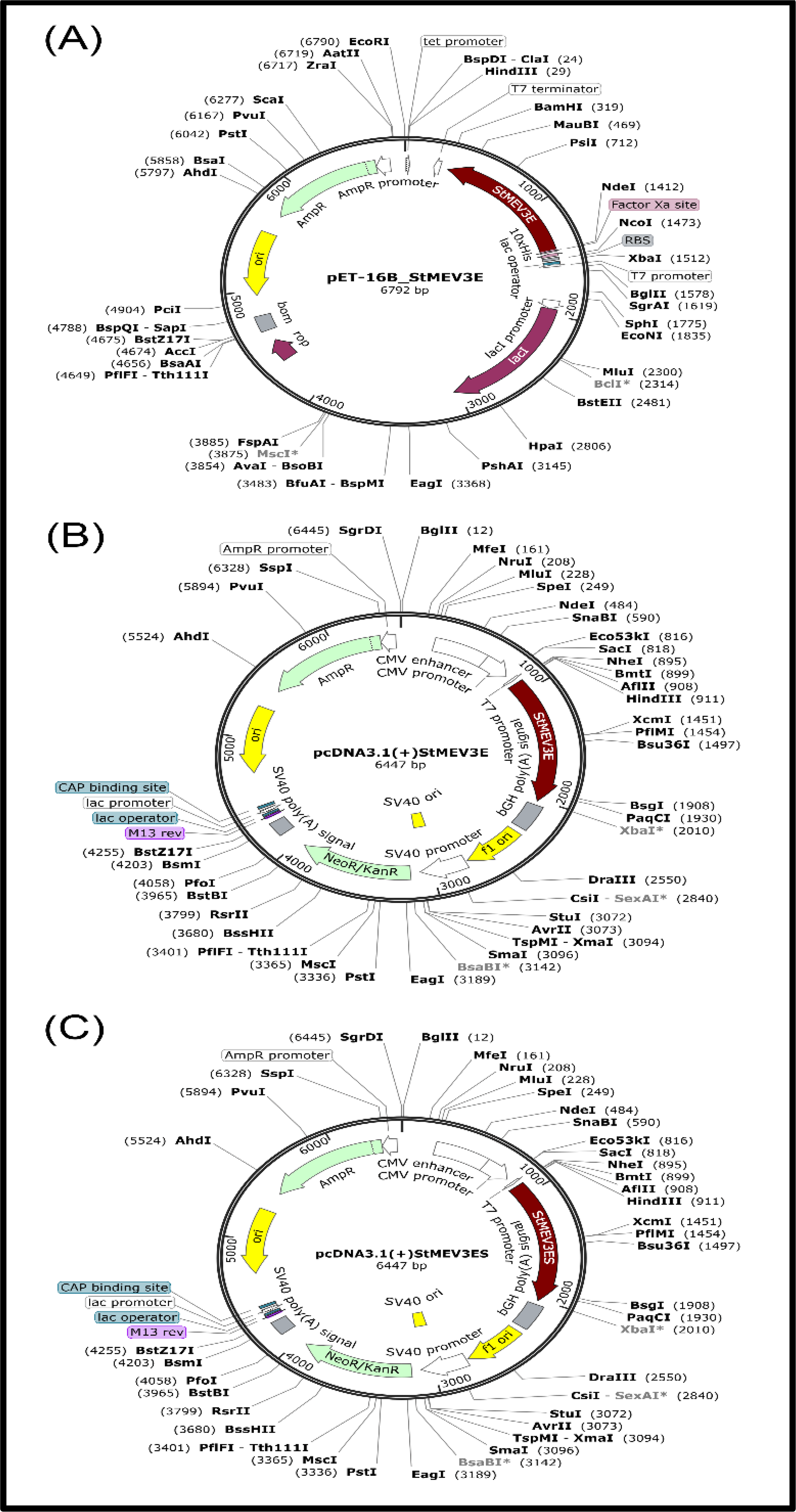
*In silico* restriction cloning and expression of STMEV3E. Codon optimized (*E. coli*) ORF of STMEV3E was cloned with in *Nde*1 and *Bam*H1 sites of pET16b to generate the construct pET16bSTMEV3E (A). Codon optimized (*H. sapiens*) ORF of STMEV3E was cloned with restriction sites for *Hind*III and *Xba*I of pcDNA3.1 (B). Codon optimized (*H. sapiens*) upstream secretory signal containing ORF of STMEV3E was cloned with restriction sites for *Hind*III and *Xba*I of pcDNA3.1.

**Fig. 5.**
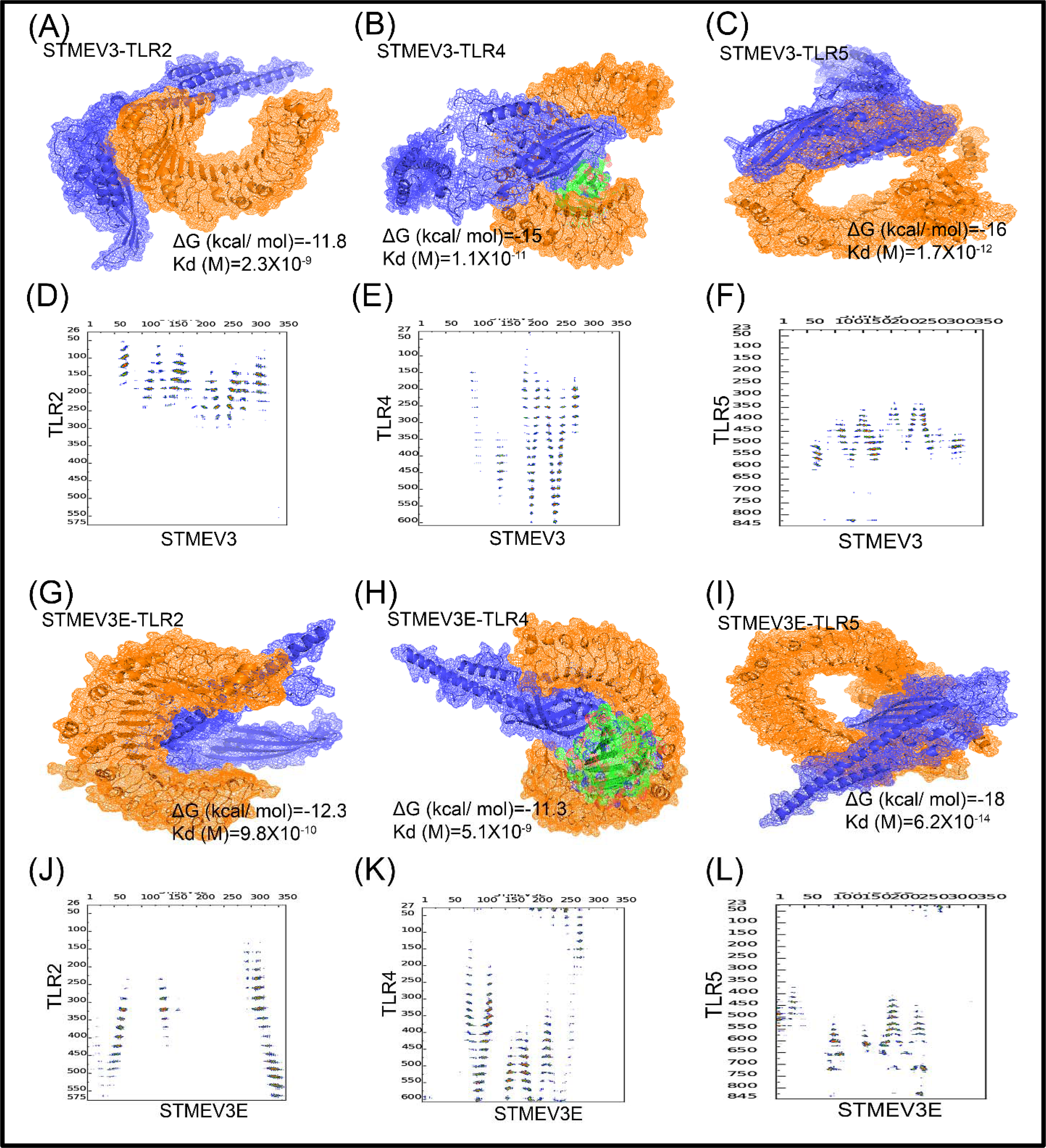
Interactions of STMEV3 and STMEV3E with TLRs. Protein-protein docking was performed by Hdock between MEVs (blue) and TLRs (orange). The predicted structures of STMEV3 was docked with available structures of TLR2 (A), TLR4 (B), and TLR5 (C). The predicted interfaces of the interactions are shown in mesh and stabilization energies (ΔG) are mentioned separately. Residue pairs involved in STMEV3-TLR2 (D), STMEV3-TLR4 (E), and STMEV3-TLR5 (F) are depicted by distance map generated through CoCoMaps. Protein-protein docking was also performed by Hdock between STMEV3E (blue) and TLRs (orange). The predicted structures of STMEV3E was docked with available structures of TLR2 (G), TLR4 (H), and TLR5 (I). The predicted interfaces of the interactions are shown in mesh and stabilization energies (ΔG) are mentioned separately. Residue pairs involved in STMEV3E-TLR2 (J), STMEV3E-TLR4 (K), and STMEV3E-TLR5 (L) are depicted by distance map generated through CoCoMaps.

The MEV-TLR complexes were further divulged for stability and dynamic behaviour by molecular dynamic simulation for 100 ns. For each of the complexes, root-mean-square deviation (RMSD), root-mean-square fluctuation (RMSF), radius of gyration (Rg), and stable H-bonds were computed for the simulation period.

RMSD of the backbone atoms during the course of simulation offers an idea about stability attainment for the complexes. Here RMSD of STMEV3 and STMEV3E were analysed for interaction with surface TLRs, TLR2, TLR4, and TLR5. For interaction with TLR2, the bound STMEV3 reached stability within 20 ns of simulation at an RMSD of 0.8 nm compared to free STMEV3, which attained stability around 70 ns at 1.5 nm (Fig. 6A). For the TLR2 in complex, stability was attained around 40 ns at 0.3 nm, while the free TLR2 attained stability around 20 ns at 0.2 nm (Fig. 6B). For STMEV3E, in complex with TLR2 reached stable RMSD of 1.00 nm around 40 ns, in contrast, free STMEV3E demonstrated less stability with RMSD of 1.6 nm around 60 ns (Fig. 6C). STMEV3E bound TLR2 and free TLR2 displayed similar RMSD profile, with the free TLR2 demonstrated initial fluctuation. TLR4-STMEV3 interaction demonstrated fluctuation for STMEV3 in complex where STMEV3 attained stability at 1.9 nm around 50 ns against free STMEV3 which demonstrated extensive fluctuation which attained stability after 70 ns at 1.5 nm (Fig. 6D). For TLR4, the STMEV3 bound and free form displayed similar profile with both reaching stable RMSD of 0.3 nm around 40 ns (Fig. 6E). STMEV3E in complex form presented greater stability at 0.9 nm attained around 15 ns, however, the free form displayed greater fluctuation (Fig. 6G). In complex with STMEV3, TLR4 demonstrated a less stable RMSD profile (Fig. 6F) compared to TLR4 in complex with STMEV3E demonstrated greater stability with stable RMSD attained at 0.18 nm around 15 ns, while the free TLR4 demonstrated RMSD around 0.3 nm throughout the simulation (Fig. 6H). For the TLR5-STMEV3 complex, the MEV in complex attained stable RMSD of 0.75 nm around 10 ns, while the free form displayed fluctuation in the initial phase and attained stability at 1.5 nm around 75 ns (Fig. 6I). TLR5 in complex demonstrated better stability compared to free TLR5 attaining stable RMSD of 1.2 nm around 30 ns and 1.4 nm around 40 ns, respectively (Fig. 6J). STMEV3E, in complex with TLR5, demonstrated immense stability in complex with RMSD of 0.5 nm attained around 5 ns of simulation in contrast to the free form demonstrating stable RMSD of 1.5 nm attained around 60 ns of simulation (Fig. 6K). In the complex TLR5 demonstrated better stability with RMSD of 1.2 nm attained within 40 ns of simulation, while the free form, displayed stable RMSD of 0.4 nm attained around 40 ns (Fig. 6L).

**Fig. 6.**
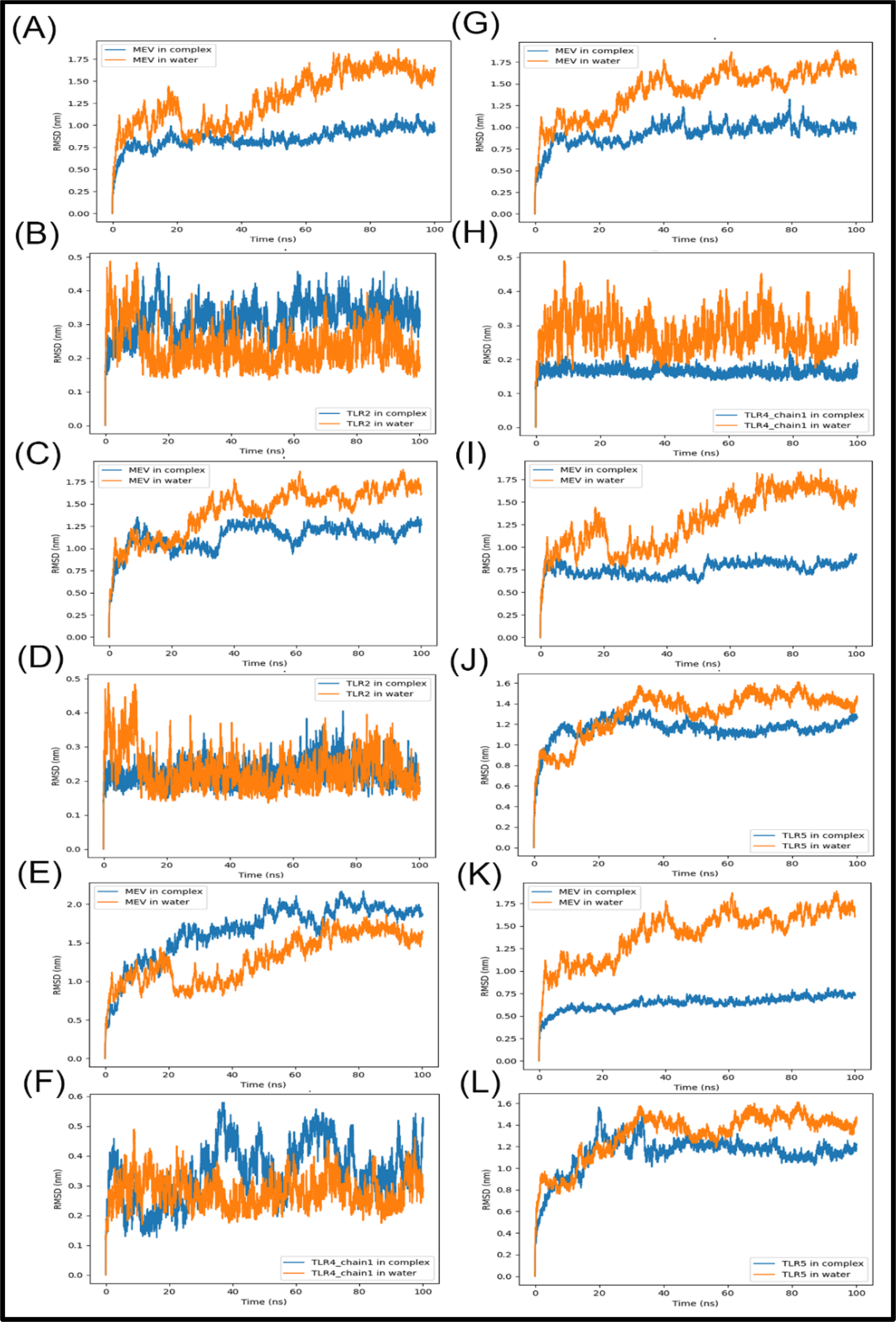
Stability attainment during MEV-TLR complex formation. RMSD of the backbone atoms during the course of simulation (100 ns) for STMEV3 (A) and TLR2 (B) in STMEV3-TLR2 complex, STMEV3E (C) and TLR2 (D) in STMEV3E-TLR2 complex, STMEV3 (E) and TLR4 (F) in STMEV3-TLR4 complex, STMEV3E (G) and TLR4 (H) in STMEV3E-TLR4 complex, STMEV3E (I) and TLR5 (J) in STMEV3-TLR5 complex, and STMEV3E (K) and TLR5 (L) in STMEV3E-TLR5 complex. Each profile was compared with respect to RMSF profile for the free form of respective MEVs or TLRs.

Structural fluctuation mediated by protein-protein interaction can be estimated by the RMSF profile, which indicates the fluctuation of amino acid residues from a reference position during the course of the simulation. For TLR2-STMEV3 interaction, the MEV in complex indicated considerable stable profile compared to the free form, with restrained fluctuation ranging between 0.18 nm to 0.75 nm for the residues as opposed to RMSF values ranging between 0.3 nm to 1.6 nm for the free form (Fig. S5A). The RMSF profile for TLR2 remained same for complex and free form with minor reduction in fluctuation for most of the residues (Fig. S5B). STMEV3E in complex with TLR2 exhibited a stable profile compared to the free MEV in water where residues positioned between 100-200 and between 300-325 demonstrated greater fluctuations reaching maximum at 0.5 nm (Fig. S5C). Similar to STMEV3-TLR2 complex, TLR2 in STMEV3E-TLR2 complex demonstrated almost identical RMSF profile (Fig. S5D). For STMEV3-TLR4 complex, STMEV3 experienced higher RMSF for majority of the N-terminal part, when in complex with TLR4; though for most of the residues between positions 225-300 it showed lesser RMSF with maximum variation ranging around 0.5 nm (Fig. S5E). For TLR4 main chain, the RMSF profile remained identical between the complex and free-form except minor variation observed in 20 residues in the C-termini (Fig. S5F). When STMEV3E form complex with TLR4, the RMSF varies for residues residing between position 60-140, 170-190, 225-250, and 275-325 compared to free form with variation ranging between 0.4-0.6 nm (Fig. S5G). Except for minor variation in the N-terminal 20 residues (∼0.1 nm), the profile remained similar for the STMEV3E bound and unbound form of the TLR4 main chain (Figs. S5H). STMEV3 in free form demonstrated greater RMDF at residue stretches 125-150, 230-250, and 260-320 with highest variation of ∼0.75 nm compared to TLR5 bound form (Fig. S5I). Considerable variation in RMSF profile was also noted for TLR5 between unbound and STMEV3-bound form with variation of RMSF between residues 690-710 and 750-800 with a maximum variation of 0.6 nm (Fig. S5J). For STMEV3E-TLR5 interaction, in complex STMEV3E demonstrated low RMSF in complex compared to the unbound form throughout with a maximum difference of ∼1 nm (Fig. S5K). RMSF profile for TLR5 also demonstrated modest variation with the complex forming chain demonstrating higher RMSF (∼1.0 nm) for the region between residues 700-780 (Fig. S5L).

To evaluate the extent of compactness attained by the MEVs during TLR interaction, radius of gyration (Rg) plots were generated. Rg plot of STMEV3 free in water indicated stabilization at 3.9 nm at 60 ns. In complex with TLR2, it attained stability around 3.9 ns at 60 ns (Fig. S6A). Rg plot for STMEV3 interaction with TLR4 indicated stabilization at 3.1 nm around 70 ns (Fig. S6B) and for interaction with TLR5 it stabilized at 3.4 nm at around 50 ns (Fig. S6C). For unbound STMEV3E, the Rg plot suggested stability at 2.7 nm at 40 ns (Fig. S6D). For interaction with TLR2, the profile indicated attaining stability at 2.6 nm at 40ns (Fig. S6E). When in complex with TLR4, the MEV attained stability at 3.0 at 10 ns and for interaction with TLR5 Rg plot indicated stability at 2.8 ns at 50 ns (Fig. S6F).

The number of stable H-bond formations is a hallmark of potential interaction between two proteins. Here for number of stable H-bond formations were simulated for STMEV3 and STMEV3E during their interaction with TLR2, TLR4, and TLR5. As depicted in Fig. S7, STMEV3 can form at least 25, 8, and 12 potential H-bonds with TLR2, TLR4, and TLR5 which were stably retained from 40 ns, 30ns, and 60 ns to 100 ns of the simulation respectively (Figs. S7A, S7B, and S7C). STMEV3E can form 18, 17, and 22 stable H-bonds which sustained from 30 ns, 50 ns, and 60 ns of the simulation to 100 ns of the simulation (Figs. S7D, S7E, and S7F).

### Immune simulation

In order to assess the impact of the designed MEVs on immunity, immune simulation was performed using C-ImmSim Immune simulator web server with default HLA setting. Two 300-day simulations were performed for three injections at 30 days intervals for 1000 STMEV3 and STMEV3E in combination with 100 adjuvants under noninfected conditions. As depicted in Figs. 7A and 7G, following the first dose, the primary response was reflected with a modest increase of IgM. Ig-levels attained remarkable and gradual elevation of IgG1, IgG1 + IgG2, IgM, and IgG + IgM antibodies following the second dose and third dose. The level of IgM + IgG attained peak after the third injection to around 2,20,000 and 2,30,000 for STMEV3 and STMEV3E respectively. Immediate depletion of the antigen after the 2^nd^ administration indicated proper clearance and induction of immunity after the1^st^administration. B-cell population and especially the B-memory cell population (∼150, ∼450, and ∼570 cells per mm^3^ for both) also increased and peaked after the third dose and the B-isotypeIgM level remained table around ∼400 cells per mm^3^ throughout the span of simulation post 1^st^ administration with small peaks around 500 cells per mm^3^ after 2^nd^ and 3^rd^ dose for both the MEVs (Figs. 7B and 7H). For STMEV3 and STMEV3E, the cytotoxic T-lymphocytes the level of active cells got elevated to ∼400 and ∼500 cells per mm^3^ and subsequently depleted (Figs. 7C and 7I). T_h_ cell population size increased after 1^st^ administration and T_h-_memory development lasted for many months between 500 to 1000 cells per mm^3^ (Figs. 7D and 7J). For NK-cells, a stable population size was registered throughout the simulation (Figs. 7E and 7K). Among the cytokines, IL-2 level was elevated around 8,00,000 ng/ ml and 8,50,000 ng/ ml for STMEV3 and STMEV3E respectively, and remained quite high (∼500000 ng/ ml) after the 3^rd^ dose, suggesting a strong humoral immune response (Figs. 7F and 7L). The role of IFN-γ production in response to *Ot* for immune protection against infection has been revealed by several studies. For STMEV3E the level of IFN-γ raised after 1^st^ and 2^nd^ doses up to 400000 ng/ ml while for STMEV3 it attained such level even after the 3^rd^ administration (Figs. 7F and 7L, inset). The projection corroborated with the identification of at least five nonoverlapping IFN-γ epitopes in STMEV3 by IFN epitope (Fig. 3J, Table-S8). Small peaks for TGF-β, Il-4, and IL-12 were also noted after each administration for both the MEVs (Figs. 7F and 7L). Such cytokine response with a low Simpson index (Figs. 7F and 7L, inset), which indicates less diversity in immune stimulation, projects potent activation of the humoral immune response.

**Fig. 7.**
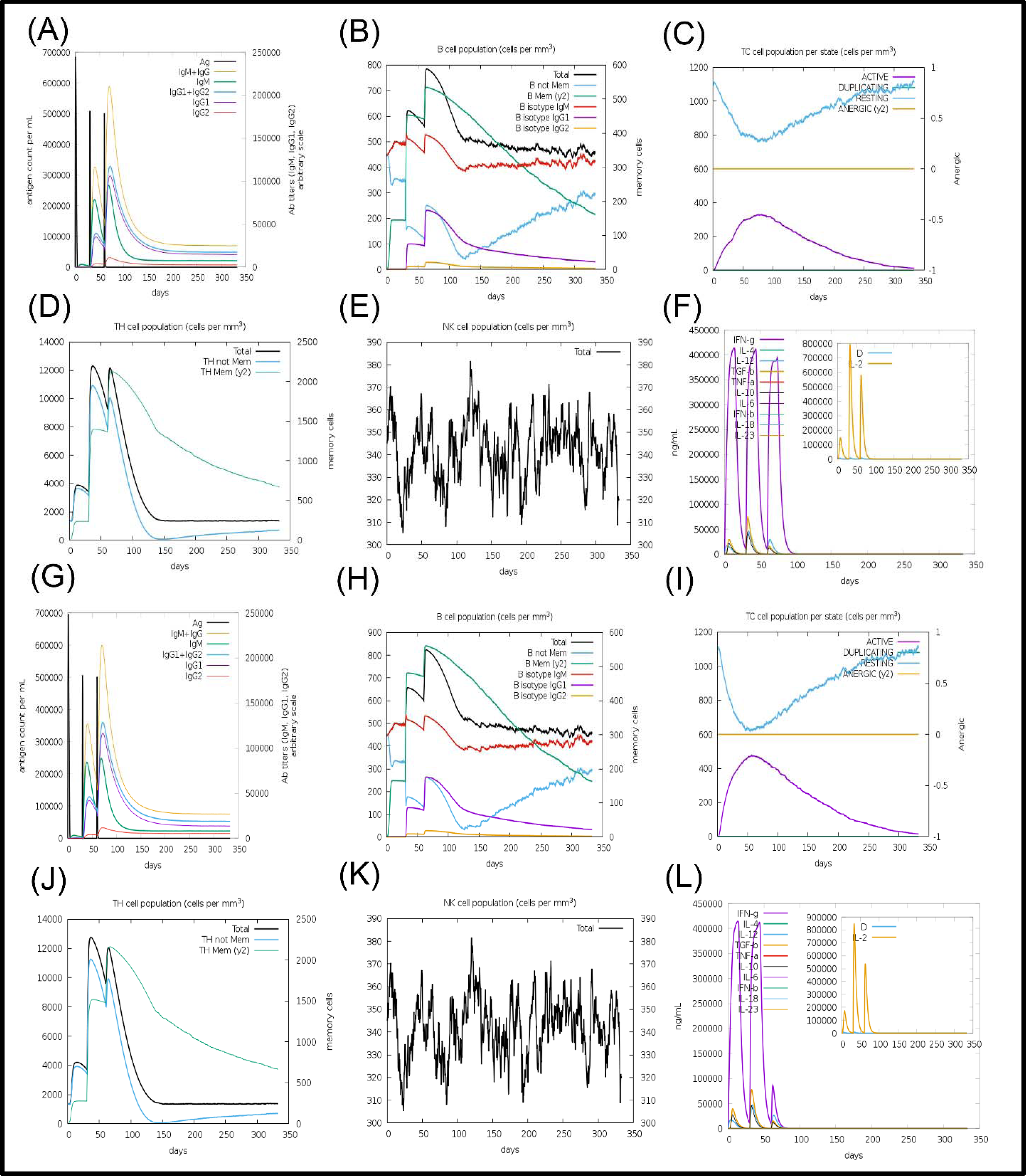
Immunosimulation for STMEV3 and STMEV3E. Immunosimulation was performed with C-ImmSim for three injections at 30 days interval. Antigen-antibody titre (A), B-cell population dynamics (B), Tc-cell population states (C), Th-cell population dynamics (D), Status of NK-cell population (E), and cytokine profile with IL-2 and Simpson index (inset) (F) are projected for each of the three doses of STMEV3. Antigen-antibody titre (G), B-cell population dynamics (H), Tc-cell population states (I), Th-cell population dynamics (J), Status of NK-cell population (K), and cytokine profile with IL-2 and Simpson index (inset) (L) are projected for each of the three doses of STMEV3E.

## Discussion

*Ot* is an obligatory intracellular Gram□negative bacterial pathogen. Developing effective vaccines against intracellular bacterial pathogens often requires activation of cytotoxic CD8+ T-cellmediated response, which is not efficiently elicited by killed whole cell- or subunit vaccines^49^. Additionally, in contrast to other intracellular bacteria like *Chlamydia* which employ lysis or extrusion for exit from host cells, *Ot* exploits an exclusive budding mechanism to become encased within host plasma membrane without affecting the integrity of host cell membrane ^50^ and thus remain unexposed to host immune system while infecting subsequent cells^51^. However, the stability of the membrane and precise organization of the trilayered membrane system remains unexplored except for the observation that along with host membrane lipid rafts, bacterial protein HtrA was detectable on the membrane ^52^. Apart from its intracellular multiplication strategy, the bacterium is considered to be fundamentally exceptional in terms of possessing canonical pathogen□associated molecular patterns (PAMP) like peptidoglycan or lipopolysaccharide^50^, though sensitivity to peptidoglycan biosynthesis targeting antibiotics ^53^and expression of genes liked to peptidoglycan biosynthesis were evidenced earlier^54^. Considering such facts here a multipronged strategy was adopted to generate a candidate MEV with the potential to elicit strong Tc-as well as Th-cell response. Strong host immune response against the bacterial proteins have been revealed earlier and the pathogen was classified into various serotypes like Gilliam, Karp, Kato, Boryong depending on the antigenic variations of the 56 kDa major surface protein, which is also the most abundant surface protein ^55^. In recent years, at least five protein antigens of molecular mass 110, 56, 47, 35, and 22 kDa were identified by immunoblotting *Ot* lysate and the type-specific antigens (TSA) 56 kDa, 47 kDa, and 22 kDa were identified as major surface protein involved in host cell adhesion ^55^. Though such tissue-specific antigens are profoundly immunodominant, and have been the obvious choice for subunit vaccine designing^56^, substantial heterogeneity of the sequence of the proteins, particularly for TSA56, impedes the broader application of such vaccine due to depreciated cross-protectivity^57^. In recent time, elicitation of protective immune response has been reported for outer membrane autotransporter proteins ScaA and ScaE, which are present in five different stains ^58^. In mouse model, ScaA inflicted protective immune response and cross protectivity against heterologous strains ^12^. Implementation of reverse vaccinology for identifying potential immunodominant proteins and sequences was augmented by whole genome sequence of the ∼2.1 mb genomes of *Ot* strain Ikeda^59^and strain Boryong ^60^ comprising 1967 and 2216 genes respectively. In 2018 long-read sequencing of five prevalent strains Karp, Kato, Gilliam, TA686, UT76 and UT176 were performed ^61^. Availability of such robust genomic data allowed us to aim at configuring MEVs through genome-wide analysis. Prediction for localization analysis allowed the identification of candidate outer membrane proteins. Interestingly, the prediction did not projected TSA56, TSA47, or TSA22 as OM proteins. When immunogenicity of the predicted OM proteins was estimated, protective immunogenicity score between 0.29 and 0.73 were detected. Considering immunogenicity for the proteins already evidenced to eliciteTSA56 (0.77), TSA47 (0.5), or TSA22 (0.57), a score of 0.5 was considered as cut-off and six OM proteins were prioritized for further analysis. Of the six OM proteins three were detected to be expressed by tanscriptomic analysis earlier and two of the three were also detected by proteomics analysis, indicating considerably high expression for the proteins ^54^. Considering the need for elicitation of both Th- and Tc-cell response, stretches that are highly enriched in sequential B-cell epitopes, Th-cell epitopes, and Tc-cell epitopes were mapped and analysed for conservancy among available genome sequences of 11 strains of *Ot*. Considering sequence diversity of apparentimmunodominat surface proteins compromise cross-protection significantly^57^, a cluster analysis was performed based on bit score obtained through BLASTP, and thus the highest ranked immmunodominant conserved segments were enlisted.

Tick salivary glands are a rich source of pharmaco-active molecules, secreted in their saliva, with immunomodulatory, anti-inflammatory, and anti-clotting activity often facilitating the tick-borne pathogens to overcome host primary defence^62^. “Acquired tick resistance” or “tick immunity,” is crucial for preventing transmission and is mediated by localized inflammatory response against tick salivary proteins. In recent times mRNA vaccines encoding salivary proteins of demonstrated to diminish *I. scapularis* engorgement on guinea pig skin and also limit *B. burgdorferi*infection ^24^. A number of immunolodulator salivary proteins have been identified from the tick with potential immunomodulatory and adjuvant action^24^. Recently 180 Mb draft genome assembly for *D. tinctorium* and a lower-coverage draft genome assembly (117 Mb) of *L. deliense*, were accomplished with the identification of orthologues of *I. scapularis* salivary proteins^25^. In this study, the salivary *I. scapularis* proteins were examined for immunogenicity and large ribosomal subunit protein P2 (A0A443SE21, *Overall Protective Antigen Prediction* score: 0.8231) and ATPase inhibitor-like protein (A0A443SL85, *Overall Protective Antigen Prediction* score: 0.7546) were identified as potentially immunogenic. Since for the large ribosomal subunit protein P2, a highly similar human orthologue was identified, immunodiminant stretch in ATPase inhibitor-like protein was identified.

Unlike the usual approach of assembling independent B-cell and T-cell epitopes to construct an immunogenic polypeptide, here epitope-enriched segments were mapped from predicted immunodominant proteins, and such stretches were assembled for configuring a combinatorial MEV. Collectively, 15 independent combinations of the identified stretches were generated and prediction for immunogenicity, allergenicity, toxicity, stability, solubility, and other physicochemical parameters allowed us to rank STMEV3 as the top-ranked MEV. Scrub typhus is a major public health problem primarily restricted in the Asia-Pacific region. The efficacy of a vaccine in a population is dependent on HLA genotypic frequency and the population coverage for a candidate vaccine is predicted based on the possibility of a selected set of epitopes with known MHC restriction. Though a population coverage emphasizing on region selective HLA allele frequency data would have offered a more intricate induction regarding the application of STMEV3, due to unavailability of such data while conducting the analysis, here, the analysis was conducted on world population coverage based on a compilation of set of alleles to cover most of the population^33^. With >95% population coverage for class I and class II MHC alleles the analysis projected STMEV3 as a potentially effective and globally administrable MEV. Glycan masking of epitopes through glycosylation of the immunogenic surface often occludes recognition by B-cell receptors^46^. In the N-termini of STMEV3, a number of potential bacterial N-linked glycosylation sites were predicted. To enhance the efficacy of STMEV3, an edited version (STMEV3E) with the glycosylable Asn (N) replaced by Gln (Q) were generated. The substitutions did not perturb, immunogenic, allergenic, and other physicochemical parameters of the MEVs and both the MEVs possess potential conformational B-cell epitopes.

TLR are receptors for proteins and other molecules including PAMPs involved in the activation of innate immune response. These receptors are localized on the cell surface (TLR1, TLR2, TLR4, and TLR5) or in intracellular compartments, such as the endoplasmic reticulum, endosome, lysosome, or endolysosome (TLR3, TLR7, TLR8, and TLR9)^48^. Strong and stable interaction of a subunit vaccine with one or more TLRs involved in dendritic cell activation, antigen processing and presentation to T_h_-cells^47^. TLR2 and 4 are also expressed in Th-cells where TLR4 signalling triggers cell proliferation, elevated IL2 and IFNγ production and subsequent surge in antibody production ^48^. In this study, possible interactions between TLR2, 4, and 5 with STMEV3 and STMEV3E were profiled by protein-protein docking. The results indicated potential interaction between the TLRs and MEVs. The binding energies ranged between -11.3 to -18.0 kcal/ mol with low K_d_ values, and considerable numbers of contact points. MD-simulation indicated STMEV3E to interact with TLR2, 4 and 5 stably as indicated by RMSD, RMSF, Rg and H-bond profile while STMEV3 interacts more stably with TLR2 and TLR5 but interaction with TLR4 appeared less stable with lesser number of steady H-bonds. Immunesimulation, as predicted for STMEV3 and STMEV4 for three injections at 30 days intervals, corroborated the TLR interaction profile. With elevated Ig levels, IFN-γ, and IL-2 and concomitant increase in B-cell and Th-cell population after administration of the MEVs. The titres of Ig, cytokines, and immune cells after each administration indicated a proper booster behaviour for both STMEV3 and STMEV3E. Such results indicated that along with recently published vaccines against various viral and bacterial pathogens, the STMEV3 and STMEV3E should be experimentally evaluated following expression and *in vitro*, *ex vivo*, and *in vivo* studies using animal models. To enable such endeavours in future here three different constructs were developed for expression and purification in bacteria, expression in mammalian cells. *In silico* cloning was performed following codon optimization for *E. coli* K12 in to the vector pET16b and following codon optimization for *H. sapiens* in pcDNA3.1. A third construct was designed by incorporating secretory signal at N-terminii of the MEVs and by subsequent *in silico* cloning in pcDNA3.1. While the first construct is aimed for expression and purification of the peptide, the second and third construct might be tested as DNA vaccine for in situ expression of the protein and elicitation of Tc-cell response as well.

In this study, we have delineated an approach aiming multiple facets of eliciting protective immune response against *Ot* infection, which includes elicitation of Th- and Tc-mediated immune response through immunodominant OM proteins of the bacteria along with tick immunity through immunodominant salivary protein of the trombidid mite vector *L. deliense*. Though the *in silico* analysis predicted potentiation of immuneresponse and immunogenic memory, *in vivo* validation and further optimization of the configurations would warrant real-world application of the proposed MEVs.

## Acknowledgement

The authors acknowledge all the open-source software and server providers. The authors are thankful to Dr.Analabha Roy, Dept. of Physics, University of Burdwan for his assistance in computational aspects. AB is funded by Startup Research Grant-SRG/2020/000702 (SERB, Govt. of India), Adhoc Grant-Project ID: 2021-14059 (ICMR, Govt. of India) and SEED grant, Adamas University.

## Availability of data and material

NA

## Author contribution

Swarna Shaw and ArkaBagchi performed most of the experiments, analysed the data.

DebyaniRuj and Sudipta Paul Bhattacharya performed the initial analysis.

Arunima Biswas was involved in the MD simulation.

Arijit Bhattacharya conceptualized the work, designed experiments, analysed data and prepared the manuscript.

All approved the final draft.

## Ethical approval

The study does not involve any human and/or animal subjects or clinical isolates. No personally identifiable patient/ human subject information was disclosed to the researchers.

## Consent to participate

NA

## Consent for publication

NA

## Reference

1. Kim DM, Kim SW, Choi SH, Yun NR. Clinical and laboratory findings associated with severe scrub typhus. BMC Infect Dis. Apr 30 2010;10:108. doi:10.1186/1471-2334-10-108

2. Chakraborty S, Sarma N. Scrub Typhus: An Emerging Threat. Indian J Dermatol. Sep-Oct 2017;62(5):478–485. doi:10.4103/ijd.IJD_388_17

3. Luce-Fedrow A, Lehman ML, Kelly DJ, et al. A Review of Scrub Typhus (Orientia tsutsugamushi and Related Organisms): Then, Now, and Tomorrow. Trop Med Infect Dis. Jan 17 2018;3(1)doi:10.3390/tropicalmed3010008

4. Kelly DJ, Fuerst PA, Ching WM, Richards AL. Scrub typhus: the geographic distribution of phenotypic and genotypic variants of Orientia tsutsugamushi. Clin Infect Dis. Mar 15 2009;48 Suppl 3:S203–30. doi:10.1086/596576

5. Lv Y, Guo XG, Jin DC. Research Progress on Leptotrombidium deliense. Korean J Parasitol. Aug 2018;56(4):313–324. doi:10.3347/kjp.2018.56.4.313

6. Walker DH, Mendell NL. A scrub typhus vaccine presents a challenging unmet need. NPJ Vaccines. Feb 9 2023;8(1):11. doi:10.1038/s41541-023-00605-1

7. Bonell A, Lubell Y, Newton PN, Crump JA, Paris DH. Estimating the burden of scrub typhus: A systematic review. PLoS Negl Trop Dis. Sep 2017;11(9):e0005838. doi:10.1371/journal.pntd.0005838

8. Yang J, Luo L, Chen T, et al. Efficacy and Safety of Antibiotics for Treatment of Scrub Typhus: A Network Meta-analysis. JAMA Netw Open. Aug 3 2020;3(8):e2014487. doi:10.1001/jamanetworkopen.2020.14487

9. Lu D, Wang T, Luo Z, et al. Evaluation of the Therapeutic Effect of Antibiotics on Scrub Typhus: A Systematic Review and Network Meta-Analysis. Front Public Health. 2022;10:883945. doi:10.3389/fpubh.2022.883945

10. Chattopadhyay S, Jiang J, Chan TC, et al. Scrub typhus vaccine candidate Kp r56 induces humoral and cellular immune responses in cynomolgus monkeys. Infect Immun. Aug 2005;73(8):5039–47. doi:10.1128/IAI.73.8.5039-5047.2005

11. Ni YS, Chan TC, Chao CC, Richards AL, Dasch GA, Ching WM. Protection against scrub typhus by a plasmid vaccine encoding the 56-KD outer membrane protein antigen gene. Am J Trop Med Hyg. Nov 2005;73(5):936–41.

12. Ha NY, Sharma P, Kim G, et al. Immunization with an autotransporter protein of Orientia tsutsugamushi provides protective immunity against scrub typhus. PLoS Negl Trop Dis. Mar 2015;9(3):e0003585. doi:10.1371/journal.pntd.0003585

13. Khan M, Khan S, Ali A, et al. Immunoinformatics approaches to explore Helicobacter Pylori proteome (Virulence Factors) to design B and T cell multi-epitope subunit vaccine. Sci Rep. Sep 16 2019;9(1):13321. doi:10.1038/s41598-019-49354-z

14. Shiragannavar S, Madagi S, Hosakeri J, Barot V. In silico vaccine design against Chlamydia trachomatis infection. Netw Model Anal Health Inform Bioinform. 2020;9(1):39. doi:10.1007/s13721-020-00243-w

15. Li M, Zhu Y, Niu C, et al. Design of a multi-epitope vaccine candidate against Brucella melitensis. Sci Rep. Jun 16 2022;12(1):10146. doi:10.1038/s41598-022-14427-z

16. Xu G, Mendell NL, Liang Y, et al. Correction: CD8+ T cells provide immune protection against murine disseminated endotheliotropic Orientia tsutsugamushi infection. PLoS Negl Trop Dis. Dec 2017;11(12):e0006127. doi:10.1371/journal.pntd.0006127

17. Hauptmann M, Kolbaum J, Lilla S, et al. Protective and Pathogenic Roles of CD8+ T Lymphocytes in Murine Orientia tsutsugamushi Infection. PLoS Negl Trop Dis. Sep 2016;10(9):e0004991. doi:10.1371/journal.pntd.0004991

18. Kodama K, Yasukawa M, Kobayashi Y. Effect of rickettsial antigen-specific T cell line on the interaction of Rickettsia tsutsugamushi with macrophages. Microbiol Immunol. 1988;32(4):435–9. doi:10.1111/j.1348-0421.1988.tb01403.x

19. Shirai A, Catanzaro PJ, Phillips SM, Osterman JV. Host defenses in experimental scrub typhus: role of cellular immunity in heterologous protection. Infect Immun. Jul 1976;14(1):39–46. doi:10.1128/iai.14.1.39-46.1976

20. Dolley A, Goswami HB, Dowerah D, et al. Reverse vaccinology and immunoinformatics approach to design a chimeric epitope vaccine against Orientia tsutsugamushi. Heliyon. Jan 15 2024;10(1):e23616. doi:10.1016/j.heliyon.2023.e23616

21. Kim CG, Kim WK, Kim N, et al. Intranasal Immunization With Nanoparticles Containing an Orientia tsutsugamushi Protein Vaccine Candidate and a Polysorbitol Transporter Adjuvant Enhances Both Humoral and Cellular Immune Responses. Immune Netw. Dec 2023;23(6):e47. doi:10.4110/in.2023.23.e47

22. Fontaine A, Diouf I, Bakkali N, et al. Implication of haematophagous arthropod salivary proteins in host-vector interactions. Parasit Vectors. Sep 28 2011;4:187. doi:10.1186/1756-3305-4-187

23. Narasimhan S, Kurokawa C, Diktas H, et al. Ixodes scapularis saliva components that elicit responses associated with acquired tick-resistance. Ticks Tick Borne Dis. May 2020;11(3):101369. doi:10.1016/j.ttbdis.2019.101369

24. Sajid A, Matias J, Arora G, et al. mRNA vaccination induces tick resistance and prevents transmission of the Lyme disease agent. Sci Transl Med. Nov 17 2021;13(620):eabj9827. doi:10.1126/scitranslmed.abj9827

25. Dong X, Chaisiri K, Xia D, et al. Genomes of trombidid mites reveal novel predicted allergens and laterally transferred genes associated with secondary metabolism. Gigascience. Dec 1 2018;7(12)doi:10.1093/gigascience/giy127

26. Yu NY, Wagner JR, Laird MR, et al. PSORTb 3.0: improved protein subcellular localization prediction with refined localization subcategories and predictive capabilities for all prokaryotes. Bioinformatics. Jul 1 2010;26(13):1608–15. doi:10.1093/bioinformatics/btq249

27. Doytchinova IA, Flower DR. VaxiJen: a server for prediction of protective antigens, tumour antigens and subunit vaccines. BMC Bioinformatics. Jan 5 2007;8:4. doi:10.1186/1471-2105-8-4

28. Magnan CN, Zeller M, Kayala MA, et al. High-throughput prediction of protein antigenicity using protein microarray data. Bioinformatics. Dec 1 2010;26(23):2936–43. doi:10.1093/bioinformatics/btq551

29. Sharma N, Patiyal S, Dhall A, Pande A, Arora C, Raghava GPS. AlgPred 2.0: an improved method for predicting allergenic proteins and mapping of IgE epitopes. Brief Bioinform. Jul 20 2021;22(4)doi:10.1093/bib/bbaa294

30. Nguyen MN, Krutz NL, Limviphuvadh V, Lopata AL, Gerberick GF, Maurer-Stroh S. AllerCatPro 2.0: a web server for predicting protein allergenicity potential. Nucleic Acids Res. Jul 5 2022;50(W1):W36–W43. doi:10.1093/nar/gkac446

31. Morozov V, Rodrigues CHM, Ascher DB. CSM-Toxin: A Web-Server for Predicting Protein Toxicity. Pharmaceutics. Jan 28 2023;15(2)doi:10.3390/pharmaceutics15020431

32. Greenbaum J, Sidney J, Chung J, Brander C, Peters B, Sette A. Functional classification of class II human leukocyte antigen (HLA) molecules reveals seven different supertypes and a surprising degree of repertoire sharing across supertypes. Immunogenetics. Jun 2011;63(6):325–35. doi:10.1007/s00251-011-0513-0

33. Weiskopf D, Angelo MA, de Azeredo EL, et al. Comprehensive analysis of dengue virus-specific responses supports an HLA-linked protective role for CD8+ T cells. Proc Natl Acad Sci U S A. May 28 2013;110(22):E2046–53. doi:10.1073/pnas.1305227110

34. Kim DE, Chivian D, Baker D. Protein structure prediction and analysis using the Robetta server. Nucleic Acids Res. Jul 1 2004;32(Web Server issue):W526–31. doi:10.1093/nar/gkh468

35. Mirdita M, Schutze K, Moriwaki Y, Heo L, Ovchinnikov S, Steinegger M. ColabFold: making protein folding accessible to all. Nat Methods. Jun 2022;19(6):679–682. doi:10.1038/s41592-022-01488-1

36. Heo L, Park H, Seok C. GalaxyRefine: Protein structure refinement driven by side-chain repacking. Nucleic Acids Res. Jul 2013;41(Web Server issue):W384–8. doi:10.1093/nar/gkt458

37. Wiederstein M, Sippl MJ. ProSA-web: interactive web service for the recognition of errors in three-dimensional structures of proteins. Nucleic Acids Res. Jul 2007;35(Web Server issue):W407–10. doi:10.1093/nar/gkm290

38. Ponomarenko J, Bui HH, Li W, et al. ElliPro: a new structure-based tool for the prediction of antibody epitopes. BMC Bioinformatics. Dec 2 2008;9:514. doi:10.1186/1471-2105-9-514

39. Dhanda SK, Vir P, Raghava GP. Designing of interferon-gamma inducing MHC class-II binders. Biol Direct. Dec 5 2013;8:30. doi:10.1186/1745-6150-8-30

40. Rapin N, Lund O, Bernaschi M, Castiglione F. Computational immunology meets bioinformatics: the use of prediction tools for molecular binding in the simulation of the immune system. PLoS One. Apr 16 2010;5(4):e9862. doi:10.1371/journal.pone.0009862

41. Yan Y, Zhang D, Zhou P, Li B, Huang SY. HDOCK: a web server for protein-protein and protein-DNA/RNA docking based on a hybrid strategy. Nucleic Acids Res. Jul 3 2017;45(W1):W365–W373. doi:10.1093/nar/gkx407

42. Xue LC, Rodrigues JP, Kastritis PL, Bonvin AM, Vangone A. PRODIGY: a web server for predicting the binding affinity of protein-protein complexes. Bioinformatics. Dec 1 2016;32(23):3676–3678. doi:10.1093/bioinformatics/btw514

43. Vangone A, Spinelli R, Scarano V, Cavallo L, Oliva R. COCOMAPS: a web application to analyze and visualize contacts at the interface of biomolecular complexes. Bioinformatics. Oct 15 2011;27(20):2915–6. doi:10.1093/bioinformatics/btr484

44. Pronk S, Pall S, Schulz R, et al. GROMACS 4.5: a high-throughput and highly parallel open source molecular simulation toolkit. Bioinformatics. Apr 1 2013;29(7):845–54. doi:10.1093/bioinformatics/btt055

45. Kumar A, Rathi E, Kini SG. Computational design of a broad-spectrum multi-epitope vaccine candidate against seven strains of human coronaviruses. 3 Biotech. Sep 2022;12(9):240. doi:10.1007/s13205-022-03286-0

46. Hariharan V, Kane RS. Glycosylation as a tool for rational vaccine design. Biotechnol Bioeng. Aug 2020;117(8):2556–2570. doi:10.1002/bit.27361

47. Duan T, Du Y, Xing C, Wang HY, Wang RF. Toll-Like Receptor Signaling and Its Role in Cell-Mediated Immunity. Front Immunol. 2022;13:812774. doi:10.3389/fimmu.2022.812774

48. Kawasaki T, Kawai T. Toll-like receptor signaling pathways. Front Immunol. 2014;5:461. doi:10.3389/fimmu.2014.00461

49. Osterloh A. Vaccination against Bacterial Infections: Challenges, Progress, and New Approaches with a Focus on Intracellular Bacteria. Vaccines (Basel). May 10 2022;10(5)doi:10.3390/vaccines10050751

50. Salje J. Orientia tsutsugamushi: A neglected but fascinating obligate intracellular bacterial pathogen. PLoS Pathog. Dec 2017;13(12):e1006657. doi:10.1371/journal.ppat.1006657

51. Fromm L, Mehl J, Keller C. Orientia tsutsugamushi: A life between escapes. Microbiologyopen. Oct 2023;12(5):e1380. doi:10.1002/mbo3.1380

52. Kim MJ, Kim MK, Kang JS. Involvement of lipid rafts in the budding-like exit of Orientia tsutsugamushi. Microb Pathog. Oct 2013;63:37–43. doi:10.1016/j.micpath.2013.06.002

53. Atwal S, Giengkam S, Chaemchuen S, et al. Evidence for a peptidoglycan-like structure in Orientia tsutsugamushi. Mol Microbiol. Aug 2017;105(3):440–452. doi:10.1111/mmi.13709

54. Cho BA, Cho NH, Min CK, et al. Global gene expression profile of Orientia tsutsugamushi. Proteomics. Apr 2010;10(8):1699–715. doi:10.1002/pmic.200900633

55. Lin CC, Chou CH, Lin TC, et al. Molecular characterization of three major outer membrane proteins, TSA56, TSA47 and TSA22, in Orientia tsutsugamushi. Int J Mol Med. Jul 2012;30(1):75–84. doi:10.3892/ijmm.2012.967

56. Yu Y, Wen B, Wen B, Niu D, Chen M, Qiu L. Induction of protective immunity against scrub typhus with a 56-kilodalton recombinant antigen fused with a 47-kilodalton antigen of Orientia tsutsugamushi Karp. Am J Trop Med Hyg. Apr 2005;72(4):458–64.

57. Ramaiah A, Koralur MC, Dasch GA. Complexity of type-specific 56 kDa antigen CD4 T-cell epitopes of Orientia tsutsugamushi strains causing scrub typhus in India. PLoS One. 2018;13(4):e0196240. doi:10.1371/journal.pone.0196240

58. Ha NY, Kim Y, Choi JH, et al. Detection of antibodies against Orientia tsutsugamushi Sca proteins in scrub typhus patients and genetic variation of sca genes of different strains. Clin Vaccine Immunol. Sep 2012;19(9):1442–51. doi:10.1128/CVI.00285-12

59. Nakayama K, Yamashita A, Kurokawa K, et al. The Whole-genome sequencing of the obligate intracellular bacterium Orientia tsutsugamushi revealed massive gene amplification during reductive genome evolution. DNA Res. Aug 2008;15(4):185–99. doi:10.1093/dnares/dsn011

60. Cho NH, Kim HR, Lee JH, et al. The Orientia tsutsugamushi genome reveals massive proliferation of conjugative type IV secretion system and host-cell interaction genes. Proc Natl Acad Sci U S A. May 8 2007;104(19):7981–6. doi:10.1073/pnas.0611553104

61. Batty EM, Chaemchuen S, Blacksell S, et al. Long-read whole genome sequencing and comparative analysis of six strains of the human pathogen Orientia tsutsugamushi. PLoS Negl Trop Dis. Jun 2018;12(6):e0006566. doi:10.1371/journal.pntd.0006566

62. Aounallah H, Bensaoud C, M’Ghirbi Y, Faria F, Chmelar JI, Kotsyfakis M. Tick Salivary Compounds for Targeted Immunomodulatory Therapy. Front Immunol. 2020;11:583845. doi:10.3389/fimmu.2020.583845

