## Supplemental_Material for "Designing Combinatorial Multiepitope Vaccine Candidates against Scrub Typus by a Proteome-wide Immunoinfomatics Approach"

**Supportive material:**

Fig. S1. Work-flow for configuring the MEVs.

Fig. S2. BLASTP analysis of *L. deliense* salivary proteins for identity with human orthologues.

Fig. S3. Secondary structure prediction for MEVs. STMEV3.

Fig. S4. *In silico*restriction cloning and expression of STMEV3.

Fig. S5. Structural fluctuation during MEV-TLR complex formation.

Fig. S6. Compactness of MEVs while interacting with TLRs.

Fig. S7. H-bond formation during complex formation between MEVs and TLRs.

Table S1. Prediction of B-cell epitopes.

Table S2. Prediction of Tc-cell epitopes.

Table S3. Prediction of Tc-cell epitopes.

Table S4. BASTP hits for immunodominant segments.

Table S5. Allergenicity prediction for MEVs.

Table S6. Conformational B-cell epitopes in STMEV3 and STMEV3E.

Table S7. Interactions in TLR-MEV complexes.

Table S8. IFN-γ epitopes in STMEV3.

Fig. S1

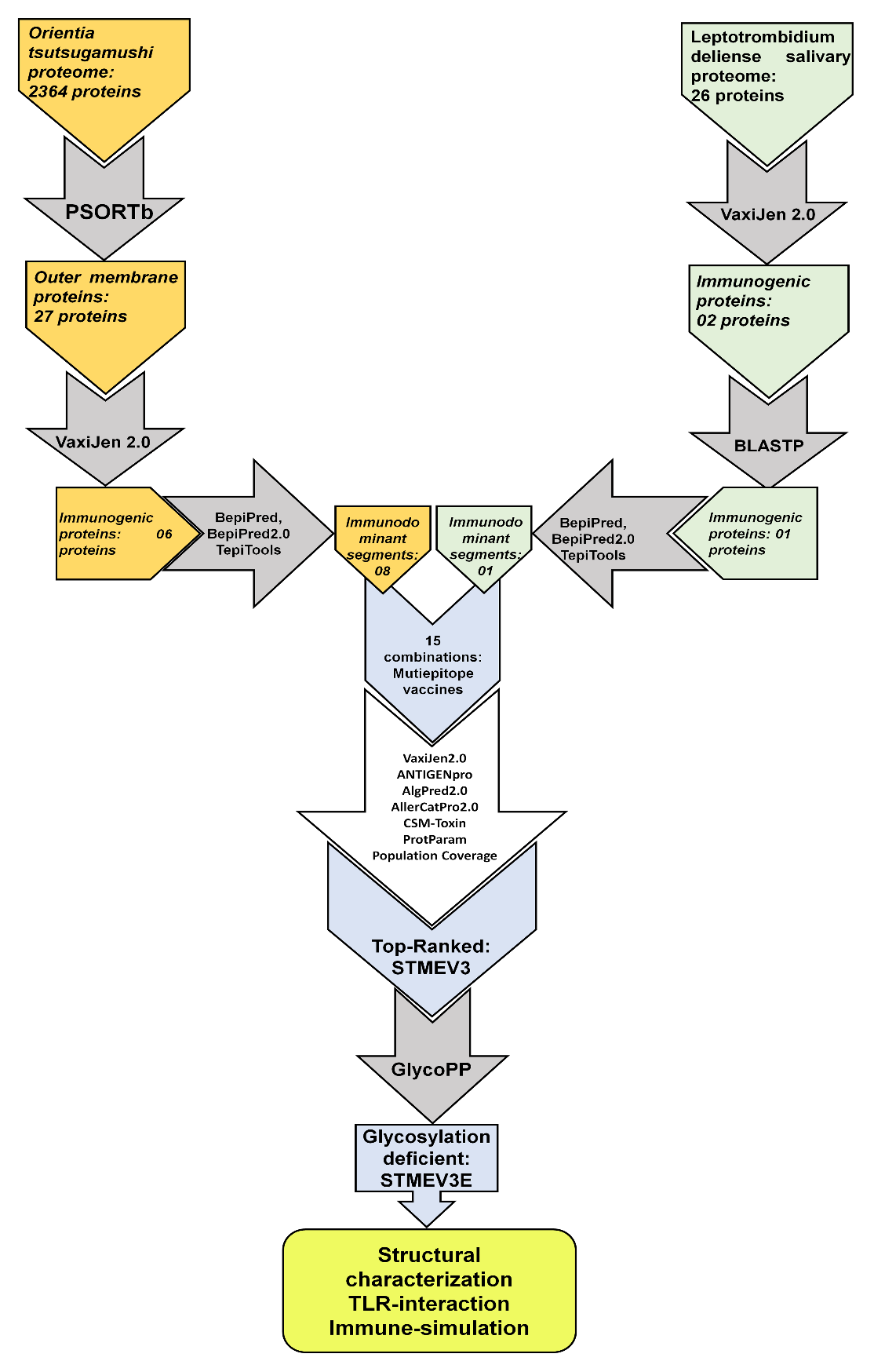

**Fig. S1.** Work-flow for configuring the MEVs. The tools implemented for each analysis are mentioned.

Fig. S2.

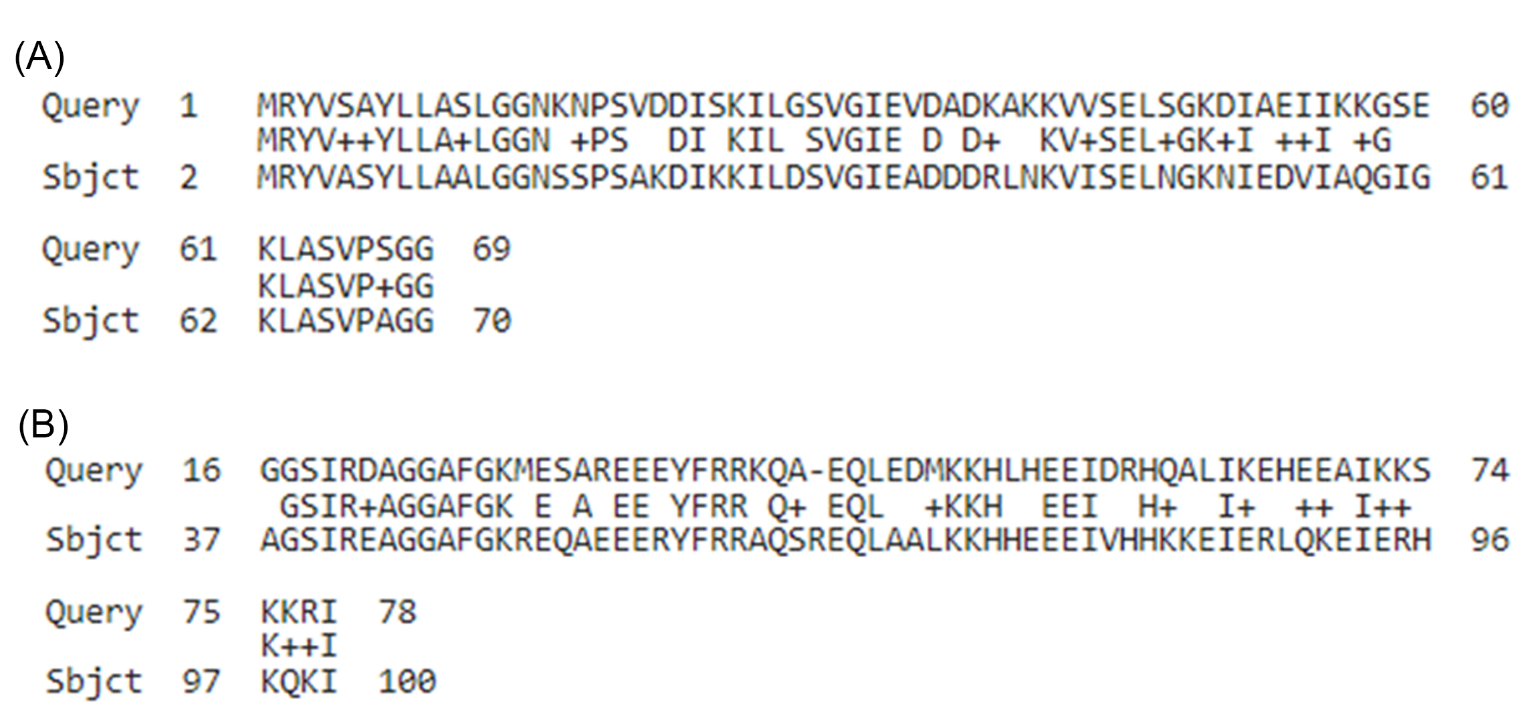

**Fig. S2.** BLASTP analysis of *L. deliense* salivary proteins for identity with human orthologues. (A) Ribosomal subunit protein P2 (A0A443SE21) and (B) ATPase inhibitor-like protein (A0A443SL85) were analysed for similarity with human orthologues. While A0A443SE21 displayed 100% coverage with 64% identity, A0A443SL85 demonstrated 73% coverage with 53.12% identity.

Fig. S3.

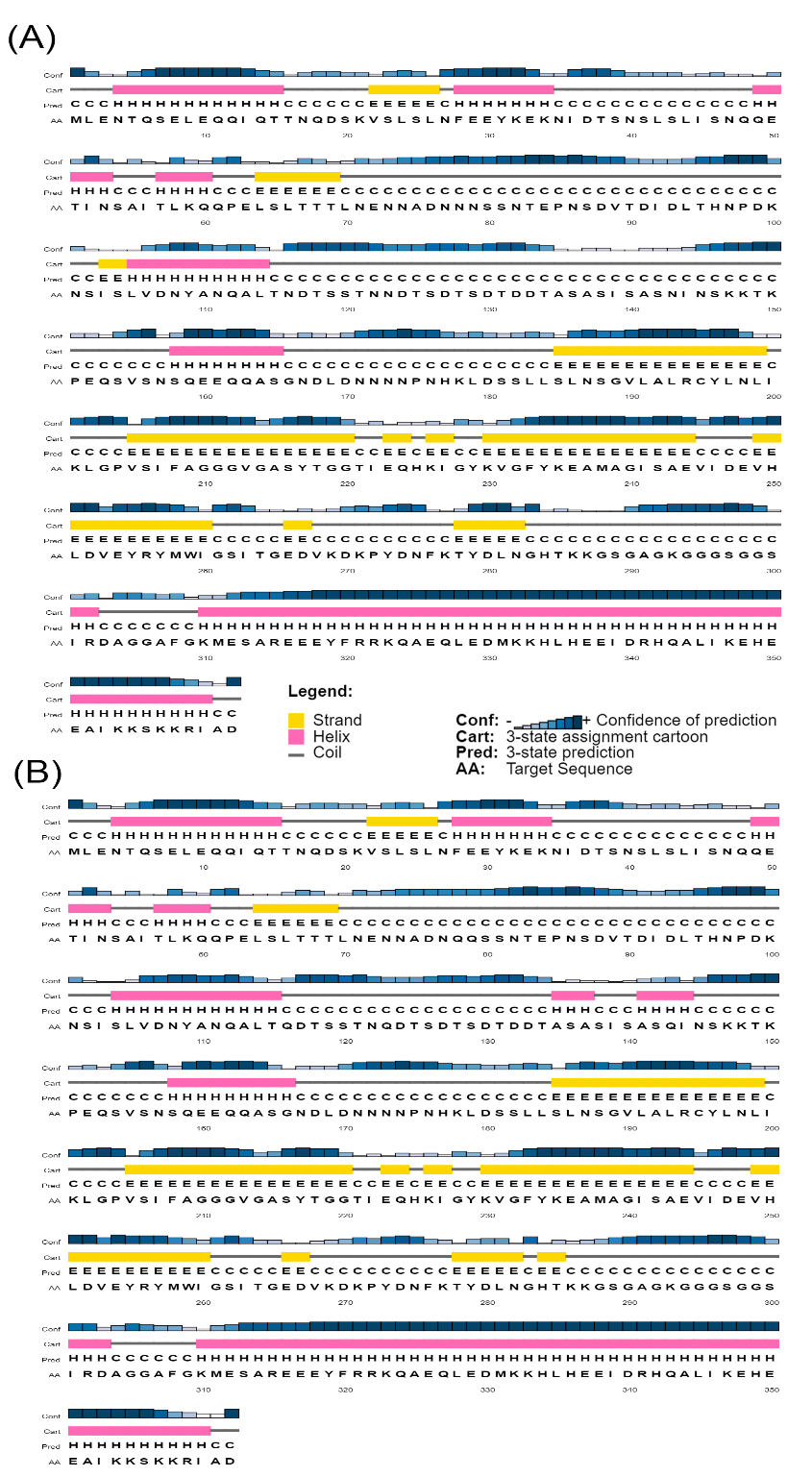

**Fig. S3.** Secondary structure prediction for MEVs. STMEV3. (A) and STMEV3E (B) sequences were analysed for secondary structure by PSI-PRED. The predicted α–helix, β-strands and coils are presented in pink block, yellow block, and blue lines respectively. Confidence of residue specific prediction is depicted by blue bars with gradient in intensity.

Fig. S4

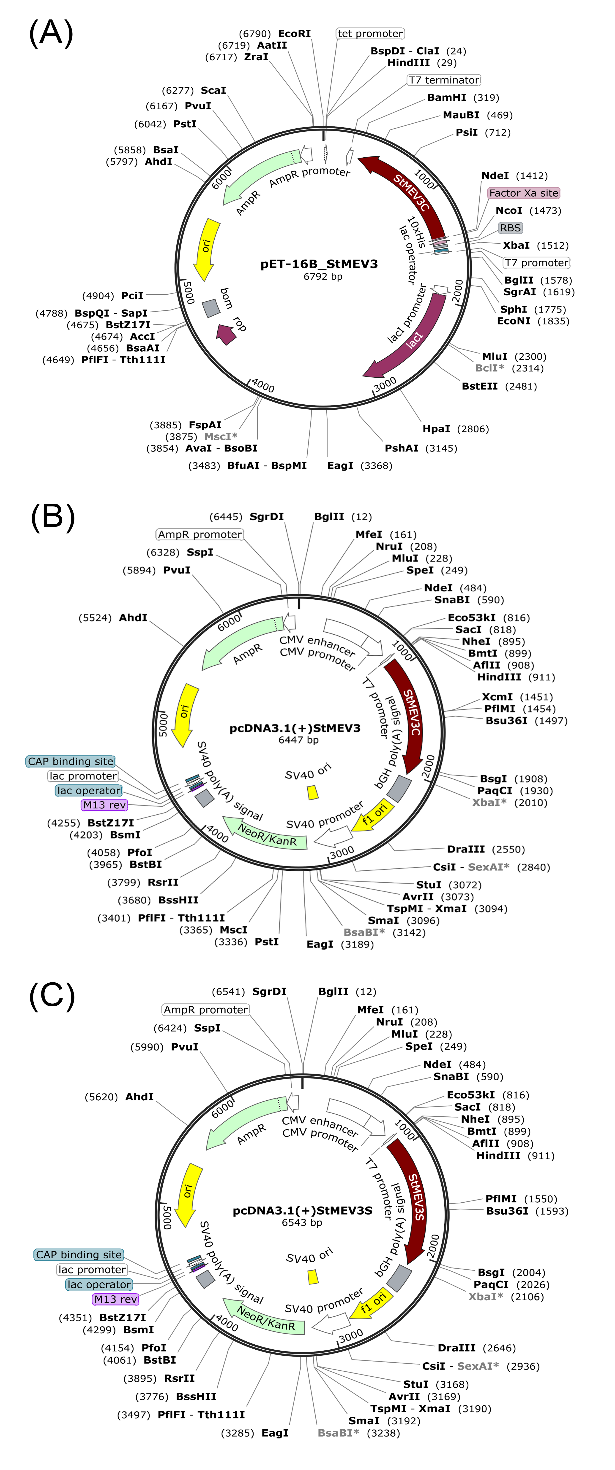

**Fig. S4.** *In silico* restriction cloning and expression of STMEV3. Codon optimized (*E. coli*) ORF of STMEV3 was cloned with in *Nde*1 and *Bam*H1 sites of pET16b to generate the construct pET16bSTMEV3 (A). Codon optimized (*H. sapiens*) ORF of STMEV3 was cloned with restriction sites for *Hind*III and *Xba*I of pcDNA3.1 (B). Codon optimized (*H. sapiens*) upstream secretory signal containing ORF of STMEV3E was cloned with restriction sites for *Hind*III and *Xba*I of pcDNA3.1.

Fig. S5

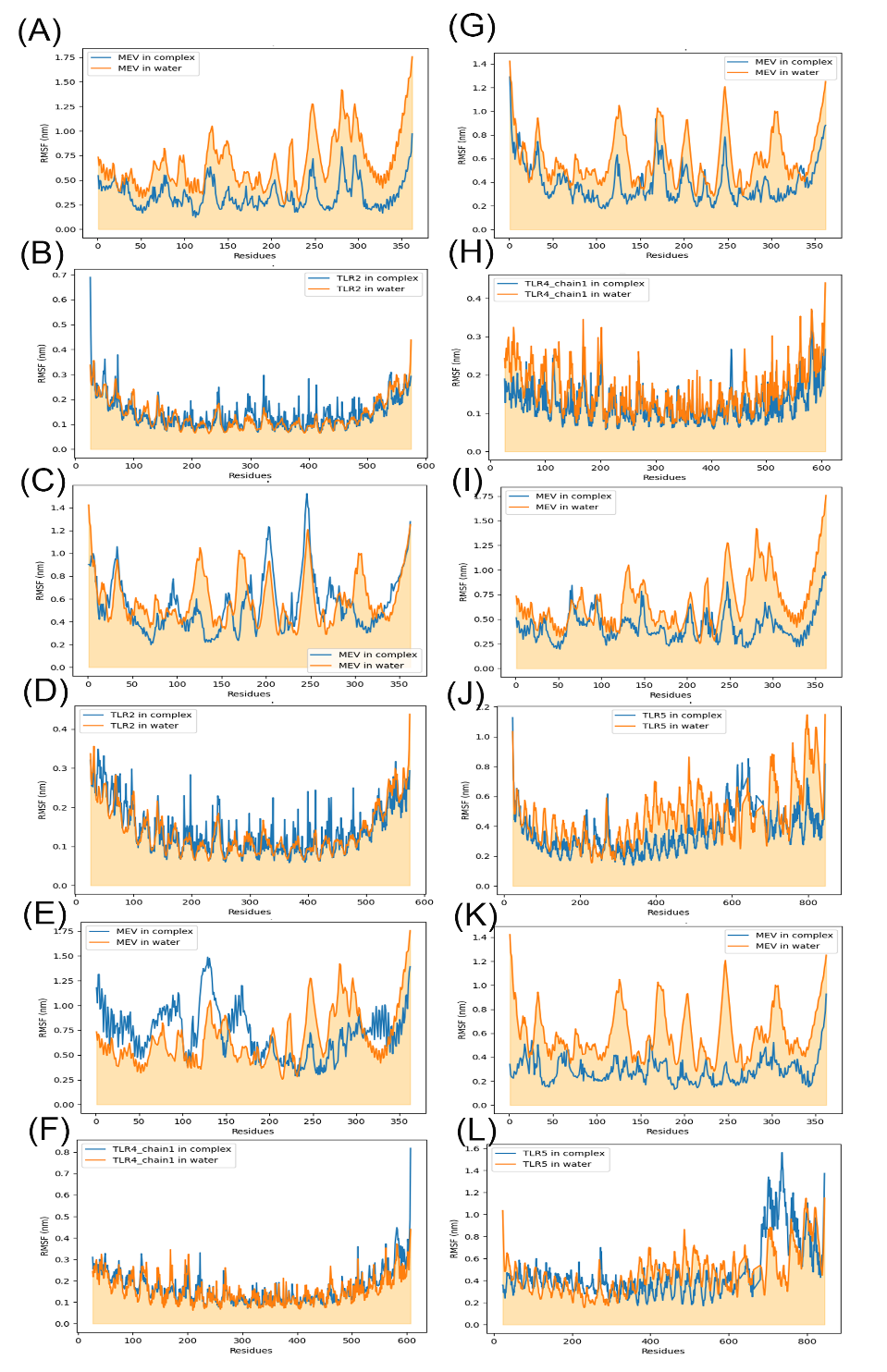

**Fig. S5.** Structural fluctuation during MEV-TLR complex formation. RMSF for amino acid residues for STMEV3 (A) and TLR2 (B) in STMEV3-TLR2 complex, STMEV3E (C) and TLR2 (D) in STMEV3E-TLR2 complex, STMEV3 (E) and TLR4 (F) in STMEV3-TLR4 complex, STMEV3E (G) and TLR4 (H) in STMEV3E-TLR4 complex, STMEV3E (I) and TLR5 (J) in STMEV3-TLR5 complex, and STMEV3E (K) and TLR5 (L) in STMEV3E-TLR5 complex. Each profile was compared with respect to RMSF profile for the free form of respective MEVs or TLRs.

Fig. S6.

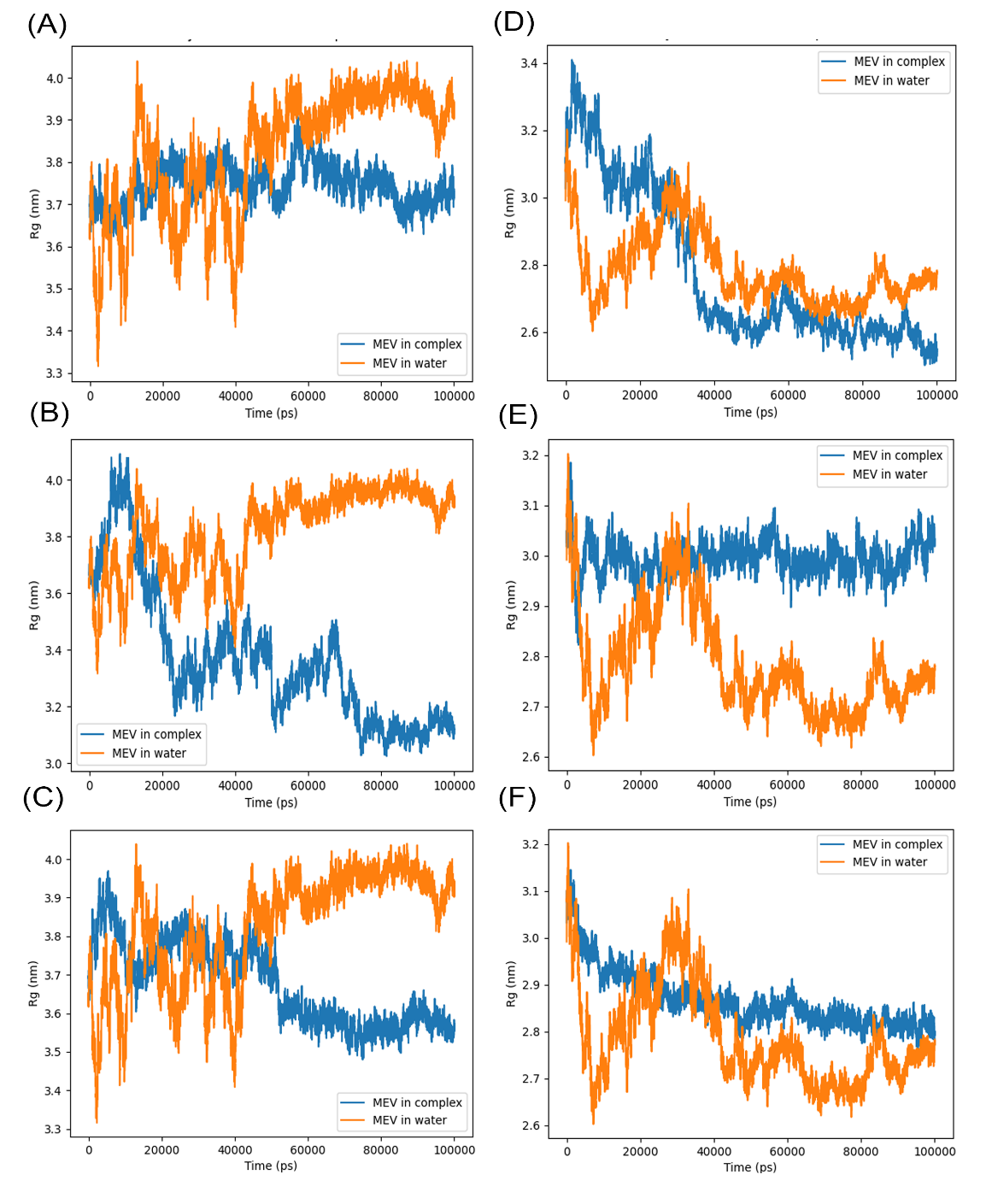

**Fig. S6.** Compactness of MEVs while interacting with TLRs. The extent of compactness for STMEV3 (A, B, and C) and STMEV3E (D, E, and F) while forming complex with TLR2 (A and D), TLR4 (B and E), and TLR5 (C and F) was envisioned by simulating radium of gyration (Rg) for 100 ns. For each complex the profile was compared with the free MEV in water.

Fig. S7.

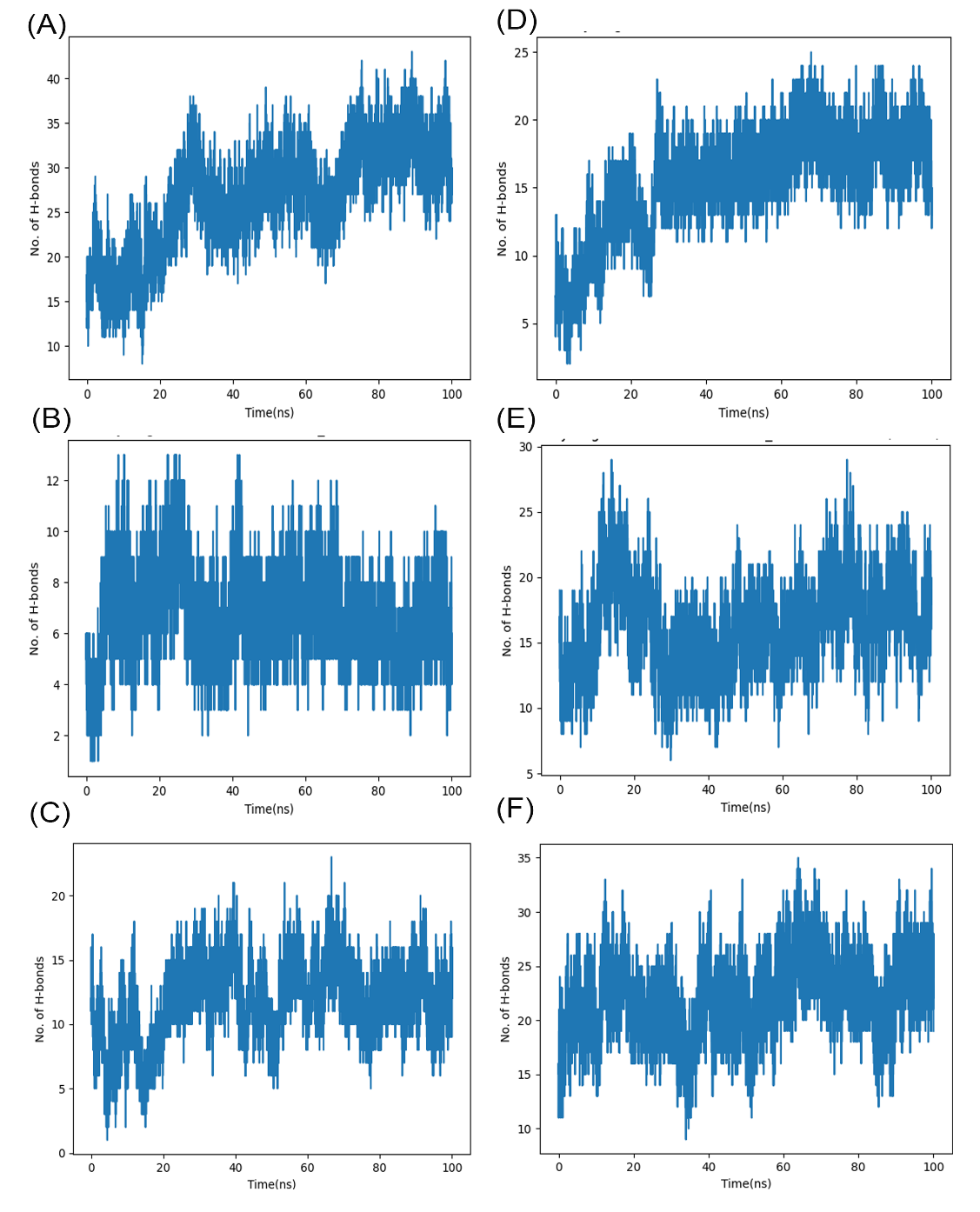

**Fig. S7.** H-bond formation during complex formation between MEVs and TLRs. The number of stable H-bond formation by STMEV3 (A, B, and C) and STMEV3E (D, E, and F) while forming complex with TLR2 (A and D), TLR4 (B and E), and TLR5 (C and F) was simulated for 100 ns.
